# A Physiologically Detailed Biomechanical Model of the Mouse Distal Forelimb for Simulation of Fine Motor Control

**DOI:** 10.64898/2026.09.14.751232

**Authors:** Nicolás Lindo-Sandoval, Jesse I. Gilmer, Géraldin Cuenu, Daniel Huber, Mazen Al Borno

## Abstract

This study presents a physiologically detailed biomechanical model of the mouse distal forelimb that incorporates intrinsic musculature, tendon routing, and digit-level skeletal anatomy, features simplified or omitted in existing musculoskeletal models. Using high-resolution anatomical reconstruction and computational modeling, we created a physiological representation of the wrist and digits capable of simulating complex forelimb movements. The model enables simulation of coordinated distal forelimb movement and digit–level muscle behavior during grasping–related tasks. Simulations were performed for multiple tasks, including grasping, grasping with supination, wrist flexion, and digit I flexion, with analysis focused on the grasping task due to its integration of both intrinsic and extrinsic musculature. Model performance was evaluated through comparisons of marker trajectories between torque–driven reference motion and muscle–driven simulations, temporal shuffle control, and comparisons between experimentally recorded electromyography (EMG) activity and model–predicted muscle excitation profiles. The model successfully reproduced coordinated distal forelimb kinematics, demonstrated strong agreement between torque-driven and muscle-driven simulation approaches, and generated physiologically plausible muscle excitation patterns consistent with experimentally observed EMG activity during grasping–related movement. These findings establish the model as a framework for studying fine motor control, neuromuscular coordination, and movement-related impairments in mice while providing a foundation for future investigation of neurological disorders and their underlying biomechanical mechanisms.

**Summary statement:** A light–sheet–derived model resolves mouse wrist and digit anatomy and tracks internally generated movements, providing a platform for experimentally testable biomechanical predictions.

## Introduction

Fine motor control of the wrist and digits plays a central role in how mice interact with their environment (Galiñanes, Bonardi and Huber, 2018). Behaviors such as skilled reaching and object manipulation require coordinated movement that begins at the shoulder, progresses through elbow extension and forearm rotation, and ultimately depends on precise control of the wrist and individual digits (Whishaw and Pellis, 1990). These skilled behaviors share important biomechanical similarities with human grasping, making the rodent distal forelimb a widely used experimental system for studying neuromuscular coordination, manipulation, and sensorimotor integration (Whishaw, Pellis and Gorny, 1992; Galiñanes, Bonardi and Huber, 2018). Consequently, characterizing the mechanics of the mouse wrist and paw is important for interpreting forelimb function in both basic neuroscience research and preclinical models of motor impairment (Sindhurakar, Butensky and Carmel, 2019).

Although interest in mouse forelimb biomechanics has grown in recent years, existing musculoskeletal models differ in their anatomical scope and reconstruction methods (DeLaurier *et al*., 2008; Ramalingasetty *et al*., 2021; Gilmer *et al*., 2024; DeWolf *et al*., 2025). Recent models have begun to incorporate wrist and digit–level structures; however, they vary in how distal skeletal elements, including the carpals and metacarpals, and muscle–tendon paths are represented (Ramalingasetty *et al*., 2021; DeWolf *et al*., 2025). In prior work, DeLaurier and colleagues mapped embryonic forelimb morphology (DeLaurier *et al*., 2008), while more recent studies, including the forelimb models of Gilmer et al. 2025 and DeWolf et al., 2025, have demonstrated the utility of anatomically informed computational models for studying forelimb biomechanics and neuromuscular function. Although the DeWolf model substantially expanded the anatomical representation of the adult mouse forelimb by incorporating wrist- and digit-level musculature, its reconstruction was based on contrast-enhanced micro-CT imaging and computational assembly in MuJoCo. Muscle–tendon paths were modeled using point-based attachments that were subsequently optimized for mechanical manipulability. This reconstruction therefore differs from the light-sheet-derived approach used here, which did not include modifying the attachment points from the imaging data. We developed the musculoskeletal model in OpenSim (Delp *et al*., 2007) rather than MuJoCo (Todorov, Erez and Tassa, 2012) as the latter implements simplified muscle modeling (e.g., by assuming inelastic tendons) for enhanced speed and simulation stability. End-users that need the computational speed (e.g., for deep reinforcement learning applications) could use our reconstruction to develop a MuJoCo model.

Although recent studies have substantially advanced the anatomical reconstruction of the mouse forelimb, evaluating how well model predictions correspond to physiological function remains a major challenge (Hicks *et al*., 2015; Gilmer *et al*., 2024; DeWolf *et al*., 2025). This challenge is particularly pronounced in the distal forelimb, where the small size and complex organization of the wrist, paw, and muscle–tendon architecture make direct experimental characterization of musculoskeletal function difficult (DeLaurier *et al*., 2008; Watson *et al*., 2009). Consequently, independent experimental measurements provide important benchmarks for assessing computational predictions of forelimb biomechanics and muscle function (Hicks *et al*., 2015).

Studying the biomechanics of the mouse’s wrist and paw is further complicated by experimental limitations, particularly in the direct measurement of muscle activity during movement. Electromyography (EMG) has been used to characterize muscle activation during mouse forelimb behaviors (Gilmer *et al*., 2024; Koh *et al*., 2025). Although EMG signals do not directly measure muscle force production or neural command, they provide information related to motor unit recruitment and neural drive to muscle, enabling qualitative comparison between experimentally recorded activity and model-predicted muscle excitation patterns (Lloyd and Besier, 2003; Buchanan *et al*., 2004; Farina, Merletti and Enoka, 2004). Previous studies have demonstrated that motor cortex output can directly influence forelimb muscle activity through corticospinal pathways, establishing a functional relationship between neural signaling and coordinated forelimb activation during movement (Cheney and Fetz, 1980; Fetz and Cheney, 1980; Lemon, 2008; Koh *et al*., 2025). Because direct experimental measurements of distal intrinsic muscle function are limited in mice, previously published EMG datasets can provide external context for examining temporal features of simulated excitation patterns. However, differences between the experimental and simulated behaviors limit direct correspondence (Hicks *et al*., 2015; Koh *et al*., 2025).

The intrinsic muscles of the paw—including the lumbricals, interossei, and deep digital flexors— are extremely small and embedded within dense musculotendinous networks, making EMG recordings and direct experimental measurements technically challenging. Additionally, the multi-articular coupling of the digits and branching of the flexor tendons make direct in vivo measurements of moment arms or digit torques in vivo challenging (Kamper, Fischer and Cruz, 2006; DeLaurier *et al*., 2008; Watson *et al*., 2009). A detailed biomechanical representation of these structures can support model–based estimates of forces produced by intrinsic and extrinsic muscles (Buchanan *et al*., 2004), and help characterize their predicted contributions to coordinated grasping (Whishaw and Pellis, 1990). An anatomically resolved musculoskeletal model of the mouse wrist and paw can therefore support simulations of joint motion, torque, and muscle excitation (Ramalingasetty *et al*., 2021; Gilmer *et al*., 2024; DeWolf *et al*., 2025).

In this study, we extended existing models of the mouse forelimb by incorporating additional distal skeletal structures, intrinsic musculature, tendon branching, and digit–level anatomy. Using anatomical segmentations derived from light-sheet microscopy, we reconstructed all carpal bones, metacarpals, phalanges, and the intrinsic and extrinsic muscles that could be resolved in the imaging dataset, including their tendon branching and insertion patterns. This model enables simulation of wrist flexion, digit I motion, grasping, and coupled pronation–supination using OpenSim Moco for musculoskeletal optimal control (Dembia *et al*., 2020). By estimating joint torques and muscle excitation across these tasks, our framework provides model–based estimates of digit–level biomechanical behavior in the mouse. For evaluation, simulated muscle excitation patterns generated during grasping with supination were compared with previously published experimentally recorded EMG data obtained during naturalistic climbing in mice, where a grasping with supination motion was being performed (Koh *et al*., 2025). This comparison was used to examine broad similarities and differences between simulated excitation profiles and experimentally observed muscle activity. Because grasping requires coordinated activation of intrinsic and extrinsic forelimb musculature, it provides a representative task for examining the temporal organization of model–predicted muscle excitation (Whishaw and Pellis, 1990; Klein *et al*., 2012). The resulting model provides an anatomically detailed computational platform for generating testable hypotheses about mouse wrist and digit biomechanics.

The objectives of this study were to (1) develop an anatomically detailed musculoskeletal model of the mouse distal forelimb incorporating intrinsic paw musculature, tendon branching, and digit– level skeletal anatomy; (2) evaluate whether muscle-driven simulations could track digit-level reference kinematics generated with torque-driven simulations; (3) quantify tracking performance across four prescribed movement tasks and a temporal-shuffling control; (4) characterize model-predicted muscle excitation during these tasks; and (5) qualitatively compare simulated muscle excitation profiles with published EMG recordings from naturalistic climbing.

## Methods

### Animals

All animal procedures used to generate the light–sheet microscopy dataset used for anatomical reconstruction were constructed as part of experiments approved by the Animal Care Committee of the University of Geneva and by the veterinary office of the “Direction Générale de la santé” of the Canton of Geneva. The anatomical reconstruction in the present study was generated from the previously acquired light–sheet microscopy dataset described by Gilmer et al. (2024), and was based on the right distal forelimb and paw of a single 11 week old female C57BL/6 mouse.

### Anatomical high-resolution imaging

Estimation of muscle activity during movement is predicated on a sufficient description of the underlying anatomy and physiology. With the goal of creating an anatomically detailed model, previously acquired anatomical data were used. High–resolution light–sheet microscopy images of the forelimb were used to reconstruct the anatomical structures included in the musculoskeletal model. These images provided the anatomical information used for segmentation and reconstruction of the skeletal and musculotendinous structures represented in the model.

### Mouse and tissue preparation

The mouse was euthanized by subcutaneous injection of pentobarbital. Transcardial perfusion with saline, followed by 4% paraformaldehyde (PFA) containing 0.01% heparin, was performed to preserve tissue integrity. The circulatory system was then washed with saline solution, followed by overnight washing with 500 mL of 1× PBS containing 0.01% heparin using transcardial perfusion and total immersion. During the washing and decalcification steps, the whole body was immersed in a chamber while the solution was circulated in a closed loop through the vascular system using a peristaltic pump. The skeletal structures were decalcified by briefly perfusing the specimen with 20% EDTA and then immersing the whole body in 500 mL of the same solution at 37°C under stirring for 31 days, with the EDTA solution renewed three times during the incubation. The specimen was subsequently washed in distilled water for 24 h. After decalcification and washing, the forelimb was dissected, the skin was removed, and the specimen was placed in a 5-mL tube for subsequent iDISCO⁺ clearing by immersion.

The dissected forelimb specimen was pretreated with methanol following the iDISCO⁺ tissue-clearing protocol. The tissue was immersed in progressively increasing concentrations of methanol, starting with 20% and increasing by 20% every hour, followed by a second 1-h immersion in 100% methanol. The initial methanol incubations were performed at room temperature. After six 1-h methanol incubations, the tissue was chilled at 4°C overnight and then immersed in 66% dichloromethane (DCM) and 33% methanol for 24 h. The tissue was then immersed twice in 100% methanol for 1 h before being chilled for 1 h at 4°C and transferred to 5% hydrogen peroxide in methanol for 48 h at 4°C without stirring. The tissue was then rehydrated by sequential immersion in 80%/60%/40%/20% methanol for 1 h per 20% decrement, then transferred to 1× PBS for 24 h, followed by a 2-h immersion at room temperature in PTx.2 (100 mL of PBS 10× and 2 mL Triton X-100, brought to a final volume of 1 L with distilled water). Given that only tissue autofluorescence was targeted for imaging, no antibodies were used in this clearing process. The tissue was permeabilized with a 500 mL solution consisting of 400 mL PTx.2, 11.5 g glycine, and 100 mL dimethyl sulfoxide (DMSO). The tissue was immersed in the solution at 37°C for 4 days, then transferred to a blocking solution containing 42 mL PTx.2 prepared as described above, 3 mL donkey serum, and 5 mL DMSO and incubated at 37°C for 3 days. The tissue was subsequently washed four times for 1 h each at room temperature with PTwH, consisting of 100 mL PBS 10×, 2 mL Tween-20, and 1 mL of 10 mg/mL heparin, brought to a final volume of 1 L with distilled water. The tissue was then re-dehydrated by sequential immersion in 20%/40%/60%/80%/100% methanol in 1 h steps, then immersed in 100% methanol overnight. Subsequently, the tissue was immersed in 66% DCM and 33% methanol for 4 h, then in 100% DCM twice for 15 min each, and finally immersed in dibenzyl ether (Sigma-Aldrich, catalog no. 108014-1KG) for imaging.

### Imaging parameters

The dissected mouse forelimb specimen was arranged in a prone position before imaging. Scans were acquired at 0.8× zoom with a pixel size of 8.23 μm and a z–step of 5 μm. Imaging was performed using a mesoSPIM (Voigt *et al*., 2019). Tissue autofluorescence was acquired in the green channel (488 nm laser) using the tiling-wizard mode with an offset of 75% and a filter set to 530/43. The forelimb was imaged in its entirety, but anatomical reconstruction was limited to the right distal forelimb and paw.

### Anatomical segmentation and reconstruction

Three-dimensional segmentation of the distal mouse forelimb was performed using the same light-sheet microscopy dataset described above. Optical slices were imported into 3D Slicer and used to manually delineate bone and muscle structures spanning the region from the elbow joint through the carpal complex and into the digits (Fedorov *et al*., 2012). Segmentation was performed manually with frequent adjustment of image brightness and contrast across axial, sagittal, and coronal views to enhance tissue boundaries. Structures were traced slice-by-slice while repeatedly verifying continuity in all three planes, using anatomical landmarks, published anatomical references, and known musculotendinous paths to guide identification of muscle bellies and tendon trajectories (DeLaurier *et al*., 2008; Watson *et al*., 2009).

Bone segmentation included the ulna, radius, a small proximal portion of the humerus (to anchor elbow geometry), all carpal bones (including the radial and ulnar carpals and carpal bones I–IV), all metacarpals, and the proximal, middle, and distal phalanges for digits II–V. Muscles of the distal compartment were segmented individually based on anatomical morphology and tendon paths, encompassing the extrinsic flexors and extensors, wrist flexors and extensors, forearm pronators and supinators, and intrinsic paw muscles (lumbricals and interossei) (DeLaurier *et al*., 2008; Watson *et al*., 2009). Following segmentation, surface meshes were exported from 3D Slicer for subsequent processing and incorporation into the musculoskeletal model. Segmentation was performed manually using 3D Slicer version 5.8.1 (Fedorov *et al*., 2012). Following completion of the segmentation, the reconstructed anatomical structures were reviewed to verify anatomical consistency before incorporating into the musculoskeletal model.

### Development of a biomechanical model of the distal mouse forelimb

Following anatomical segmentation, the distal forelimb geometry was assembled into a biomechanical framework using OpenSim Creator (Kewley, Beesel and Seth, 2026). Surface meshes generated from 3D Slicer were exported to Blender version 5.1.1 for mesh processing and bone and muscle volume calculations (Development Team, 2026). Muscle meshes were visually inspected and adjusted to maintain correspondence with the tendon paths identified in the optical slices before the processed meshes were imported into OpenSim Creator for model reconstruction (Fig. 1B, C). The global model coordinate system was defined with the X-axis oriented proximodistally, the Y-axis oriented dorsopalmarly, and the Z-axis oriented radioulnarly. Gravity was applied along the negative Y-axis. In Figure 1C, the model is displayed with the elbow flexed to approximately 50°, with the positive Y-axis pointing upward, to represent the limb in an anatomically interpretable orientation relative to gravity.

**Fig. 1.**
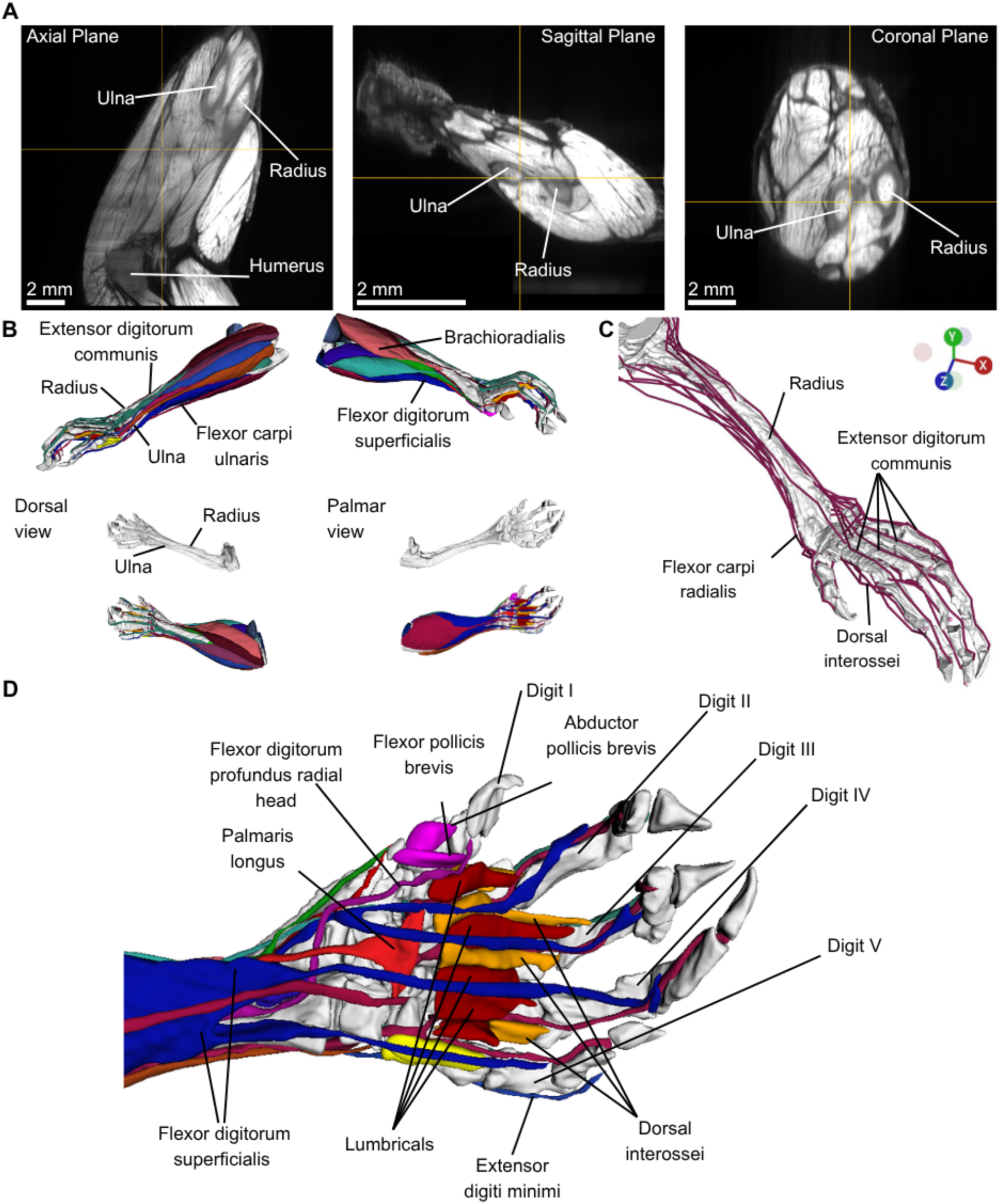
Anatomical reconstruction of the distal mouse forelimb. (A) Optical slices of the distal forelimb in the axial, sagittal, and coronal planes, shown in the prone orientation, with the radius, ulna, and humerus labeled to illustrate anatomical landmarks used during segmentation. (B) Three-dimensional reconstructions of the segmented distal skeleton and musculature displayed from multiple viewpoints. Bone-only renderings demonstrate the overall distal limb morphology, while composite reconstructions highlight the organization of major extrinsic and intrinsic muscle groups, including the extensor digitorum communis, flexor carpi ulnaris, flexor digitorum superficialis, and brachioradialis (Zhang, Zhang and Medler, 2010), illustrating the dense musculotendinous architecture of the wrist and digits. The smaller renderings show dorsal views in the left column and palmar views in the right column. (C) Composite three-dimensional reconstruction of the distal musculature mapped onto the skeletal geometry, with selected muscles labeled to illustrate their anatomical locations and routing along the forelimb. The muscles shown include the extensor digitorum profundus, extensor digitorum communis, extensor carpi radialis longus, flexor carpi radialis, flexor carpi ulnaris, abductor pollicis longus, pronator teres, and flexor digitorum superficialis. (D) Detailed three-dimensional reconstruction of the distal forelimb and paw, with digits I–V identified. Selected intrinsic and extrinsic muscles include the lumbricals, dorsal interossei, flexor pollicis brevis, flexor digitorum profundus radial head, palmaris longus, and extensor digiti minimi. These structures illustrate the dense tendon architecture and complex muscle routing that characterize the distal limb. The coordinate triad in the top-left panel denotes the global model reference frame as defined in OpenSim Creator.

Following import into OpenSim Creator, segment mass properties were defined for each bone. The center of mass for each segment was estimated by a geometric marker placed at the centroid of the corresponding mesh. Bone volumes were computed in Blender version 5.1.1. using the 3D Print Toolbox (Development Team, 2026) after applying the same linear geometric scale factor (1.91 × 10^−6^) used during mesh import. This scale factor was stablished during the original Slicer–to–Blender export workflow to convert the reconstructed geometry to the scale required for OpenSim; however, the intermediate derivation of the factor was not retained. Segment mass was calculated as *m* = *ρV*, where *ρ* = 2000 *^kg^*/*_m_*_3_ represents bone density and *V* is segment volume, consistent with previously established biomechanical modeling approaches for the mouse forelimb (Ramalingasetty *et al*., 2021). Consistent with the approach described by Gilmer et al. (2025), bone-specific inertia tensors were estimated from the corresponding reconstructed mesh geometries under the same uniform-density assumption. Each inertia tensor was calculated about the segment’s center of mass, expressed in the corresponding local body coordinate system, and assigned during the OpenSim model construction.

Bones were defined as rigid bodies and organized into a kinematic chain representing the distal forelimb from the elbow through the wrist and into the digits. Joints were created to correspond to each anatomically distinct articulation, including the wrist, metacarpophalangeal, proximal interphalangeal, and distal interphalangeal joints, as well as the digit I-specific articulations. Joint coordinate axes were explicitly defined to reflect the primary anatomical movements at each joint, and joint ranges of motion were constrained to anatomically plausible limits based on anatomical considerations and manual manipulation of each joint within OpenSim Creator. The limits were adjusted until the resulting movements were judged anatomically plausible. This was intended to maintain consistency with the reconstructed murine joint anatomy while preventing anatomically implausible movement.

Muscle paths were incorporated using multiple path points for each muscle, with all points defined in the local coordinate frames of their respective parent bones. Path points were parented to the appropriate anatomical structures to preserve the reconstructed tendon routing throughout joint motion and to maintain stable muscle–bone relationships across the full range of movement. This geometric formulation prioritized correspondence with the reconstructed anatomy and defined the muscle paths used in the subsequent simulations before dynamic and optimization settings were introduced.

### Muscle path definition and attachment locations

Muscles were represented using De Groote–Fregly 2016 Hill-type musculotendon actuators within OpenSim (De Groote *et al*., 2016). Muscle paths were defined in OpenSim Creator using the segmented anatomy as a geometric reference. For each muscle, the parent bodies for the origin and insertion were assigned by tracing the reconstructed muscle and tendon paths in the optical image slices. For muscles whose insertions occurred primarily in fascial structures rather than on a single discrete bone, the insertion was assigned to the closest anatomically relevant bone surface to approximate the functional attachment site. Path point coordinates were extracted directly from OpenSim Creator and recorded in the local coordinate frame of their respective parent bodies.

Intermediate path points were added for each muscle as needed to approximate anatomically plausible tendon routing. Path point placement and parent bone assignment were determined by evaluating joint motion and identifying which skeletal elements influenced tendon trajectory during movement. Path points were parented to the bones most closely associated with the corresponding joint motion to allow the muscle paths to change appropriately as joints moved. The number of path points assigned to each muscle was determined iteratively by manually testing joint motion and adjusting muscle paths to prevent anatomically implausible intersections or ovelap between adjacent muscles and tendons. This process was repeated until muscle paths remained separated and anatomically plausible across the full range of allowable joint motion. For muscles in which a discrete tendon could not be reliably distinguished from the reconstructed anatomy, tendon slack length was assigned a value of zero, consistent with the simplified musculotendon representation adopted in this model. Representative muscle origin and insertion locations are summarized in Table 1, while complete origin and insertion locations for all modeled muscles are provided in Table S1.

**Table 1.**
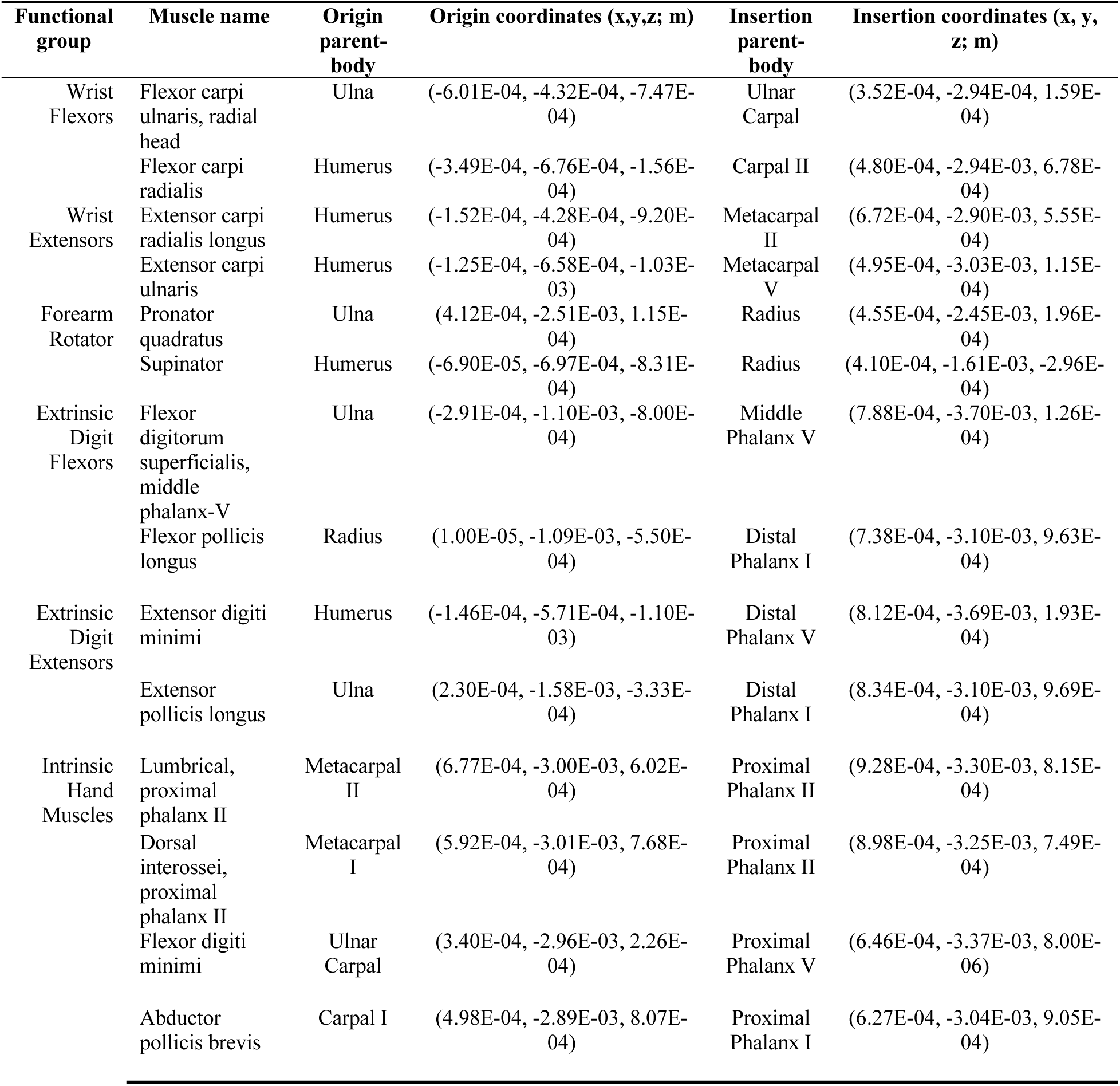
Representative muscle origin and insertion locations organized by functional group. Muscle origin and insertion locations are reported in the local coordinate frame of their parent bodies. Representative muscles spanning the major intrinsic and extrinsic functional groups of the mouse forelimb are shown for clarity. Complete muscle origin and insertion data for all modeled muscles are provided in Table S1.

| Functional group | Muscle name | Origin parent-body | Origin coordinates (x,y,z; m) | Insertion parent-body | Insertion coordinates (x, y, z; m) |
| --- | --- | --- | --- | --- | --- |
| Wrist Flexors | Flexor carpi ulnaris, radial head | Ulna | (-6.01E-04, -4.32E-04, -7.47E-04) | Ulnar Carpal | (3.52E-04, -2.94E-04, 1.59E-04) |
|  | Flexor carpi radialis | Humerus | (-3.49E-04, -6.76E-04, -1.56E-04) | Carpal II | (4.80E-04, -2.94E-03, 6.78E-04) |
| Wrist Extensors | Extensor carpi radialis longus | Humerus | (-1.52E-04, -4.28E-04, -9.20E-04) | Metacarpal II | (6.72E-04, -2.90E-03, 5.55E-04) |
|  | Extensor carpi ulnaris | Humerus | (-1.25E-04, -6.58E-04, -1.03E-03) | Metacarpal V | (4.95E-04, -3.03E-03, 1.15E-04) |
| Forearm Rotator | Pronator quadratus | Ulna | (4.12E-04, -2.51E-03, 1.15E-04) | Radius | (4.55E-04, -2.45E-03, 1.96E-04) |
|  | Supinator | Humerus | (-6.90E-05, -6.97E-04, -8.31E-04) | Radius | (4.10E-04, -1.61E-03, -2.96E-04) |
| Extrinsic Digit Flexors | Flexor digitorum superficialis, middle phalanx-V | Ulna | (-2.91E-04, -1.10E-03, -8.00E-04) | Middle Phalanx V | (7.88E-04, -3.70E-03, 1.26E-04) |
|  | Flexor pollicis longus | Radius | (1.00E-05, -1.09E-03, -5.50E-04) | Distal Phalanx I | (7.38E-04, -3.10E-03, 9.63E-04) |
| Extrinsic Digit Extensors | Extensor digiti minimi | Humerus | (-1.46E-04, -5.71E-04, -1.10E-03) | Distal Phalanx V | (8.12E-04, -3.69E-03, 1.93E-04) |
|  | Extensor pollicis longus | Ulna | (2.30E-04, -1.58E-03, -3.33E-04) | Distal Phalanx I | (8.34E-04, -3.10E-03, 9.69E-04) |
| Intrinsic Hand Muscles | Lumbrical, proximal phalanx II | Metacarpal II | (6.77E-04, -3.00E-03, 6.02E-04) | Proximal Phalanx II | (9.28E-04, -3.30E-03, 8.15E-04) |
|  | Dorsal interossei, proximal phalanx II | Metacarpal I | (5.92E-04, -3.01E-03, 7.68E-04) | Proximal Phalanx II | (8.98E-04, -3.25E-03, 7.49E-04) |
|  | Flexor digiti minimi | Ulnar Carpal | (3.40E-04, -2.96E-03, 2.26E-04) | Proximal Phalanx V | (6.46E-04, -3.37E-03, 8.00E-06) |
|  | Abductor pollicis brevis | Carpal I | (4.98E-04, -2.89E-03, 8.07E-04) | Proximal Phalanx I | (6.27E-04, -3.04E-03, 9.05E-04) |

Muscle-specific parameters were assigned based on geometric measurements and standard musculoskeletal modeling assumptions. Due to the limited availability of direct experimental measurements for mouse distal forelimb musculature, parameters such as pennation angle, tendon slack length, and maximum isometric force were estimated using values and assumptions reported in prior musculoskeletal modeling studies, while muscle geometric properties were obtained from the reconstructed anatomy. Whole-muscle geometric lengths were obtained from the reconstructed geometry and converted to physical units using a linear conversion factor of 0.00512 mm per measurement unit. These measurements represent the reconstructed whole-muscle length and should not be interpreted as muscle fiber or fascicle length, which can differ from whole-muscle length depending on muscle architecture and pennation (Roberts *et al*., 2019). This value was used only to convert measurements obtained from the reconstructed geometry and does not represent the native image voxel spacing; the image volume was not resampled. Anatomical cross-sectional area (ACSA) was estimated by dividing reconstructed muscle volume by total muscle length. This quantity differs from physiological cross-sectional area (PCSA), which is conventionally calculated using muscle fascicle length and, when included, pennation angle (van Leeuwen *et al*., 2018). Because fascicle lengths were not available for the reconstructed distal forelimb muscles, PCSA could not be directly determined. Maximum isometric force was therefore approximated by multiplying the estimated ACSA by a specific muscle tension of 28 N/cm², and these values should be interpreted as geometry-based estimates rather than experimentally derived physiological force capacities. For muscles represented by multiple heads or paths, the force calculated for each reconstructed muscle component was divided by the number of corresponding heads or paths. Because force was calculated independently for each reconstructed muscle component using its measured geometry, paired heads received unequal force values. For muscles in which a distinct free-tendon segment was not represented or could not be reliably measured from the reconstructed anatomy, including pronator teres, pronator quadratus, supinator, and several intrinsic hand muscles, tendon slack length was assigned a value of zero. Complete muscle-specific parameter values for all modeled muscles are provided in Table S2.

### Joint coordinate definitions and model degrees of freedom

The musculoskeletal model contained 28 joints, including 18 mobilized joints that provided 24 rotational degrees of freedom and 10 welded joints. The humerus was fixed to ground, the ulnar carpal and carpal I–IV segments were welded to the radial carpal segment, and the carpometacarpal joints of digits II–V were welded. Elbow flexion was represented by rotation of the ulna relative to the humerus, whereas forearm supination–pronation was represented by rotation of the radius relative to the ulna. The wrist was represented by flexion–extension and ulnar–radial deviation of the radial carpal segment relative to the radius. Digit I included four rotational coordinates: flexion–extension and abduction–adduction at the carpometacarpal joint, flexion of the proximal phalanx at the metacarpophalangeal joints, and flexion of the distal phalanx at the interphalangeal joint. Each of digits II–V also included four rotational coordinates: flexion–extension and abduction–adduction at the metacarpophalangeal joints, middle phalanx flexion at the proximal interphalangeal joint, and distal phalanx flexion at the distal interphalangeal joint. Joint limits were encoded in the biomechanical model for all degrees of freedom. The modeled joint coordinates and their anatomical motions are illustrated in Figure 2.

**Fig. 2.**
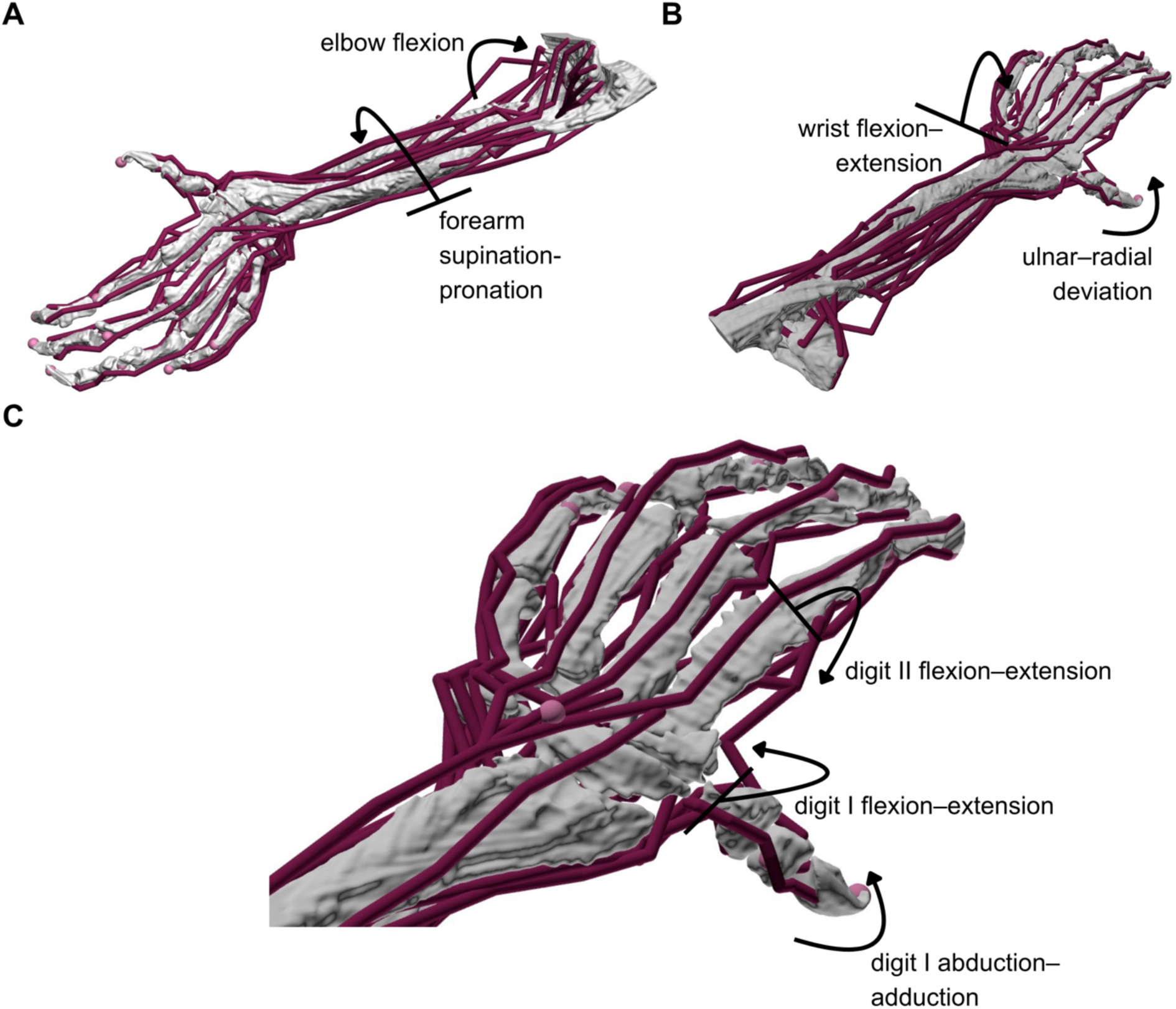
Representative joint coordinates and their corresponding anatomical motions in the musculoskeletal model. (A) Elbow flexion, represented by rotation of the ulna relative to the humerus, and forearm supination–pronation, represented by rotation of the radius relative to the ulna. (B) Wrist flexion–extension and ulnar–radial deviation of the radial carpal segment relative to the radius. (C) Digit I flexion–extension and abduction–adduction at the carpometacarpal joint and digit II flexion–extension at the metacarpophalangeal joint. The digit II coordinate is shown as representative of the corresponding flexion–extension coordinates in digits II–V. Curved arrows indicate the rotational motions associated with the selected coordinates and do not specify their positive directions. Muscle paths are shown for anatomical context.

### Kinematic and muscle-driven simulations

Simulations were performed with OpenSim 4.5 and OpenSim Moco (Dembia *et al*., 2020) using a Python environment (Python 3.13.9). We simulated four prescribed distal forelimb movements: grasping, grasping with supination, digit I flexion, and wrist flexion. Optimization was performed using the CasADi solver with a direct collocation formulation consisting of 60 mesh intervals. Convergence and constraint tolerances were set to 1 × 10⁻⁵ for the torque-driven optimization and 1 × 10⁻⁴ for the muscle-driven optimization, respectively, with a maximum of 500 optimization iterations. Each movement was simulated independently using a two-step workflow. In the first step, a torque-driven model was used to generate reference marker trajectories from prescribed initial and final joint angles. These simulations were formulated as optimal control problems using OpenSim Moco, where joint coordinate actuators replaced musculotendon units and a control effort term was defined as the sum of the squared coordinate–actuator controls. Coordinate actuator strengths and control bounds were specified within the OpenSim model (Table S3). In the second step, the resulting marker trajectories were used as tracking targets for a muscle-driven model through marker tracking. Three-dimensional marker trajectories were applied as tracking targets using a MocoMarkerTrackingGoal, with all experimental markers assigned equal weights (1 × 10⁹), while a control effort term was included to regularize the muscle excitations using a MocoControlGoal with a weight of 1. The objective function consisted of weighted marker–tracking and control–effort terms, where the control effort was defined as the sum of the squared muscle excitations. Complete optimization settings, including software versions, optimization parameters, actuator settings, control bounds, marker weights, and prescribed coordinate values, are provided in Table S3. An overview of the simulated grasping movement and the corresponding resulting joint–angle trajectories are shown in Figure 3A, B.

**Fig. 3.**
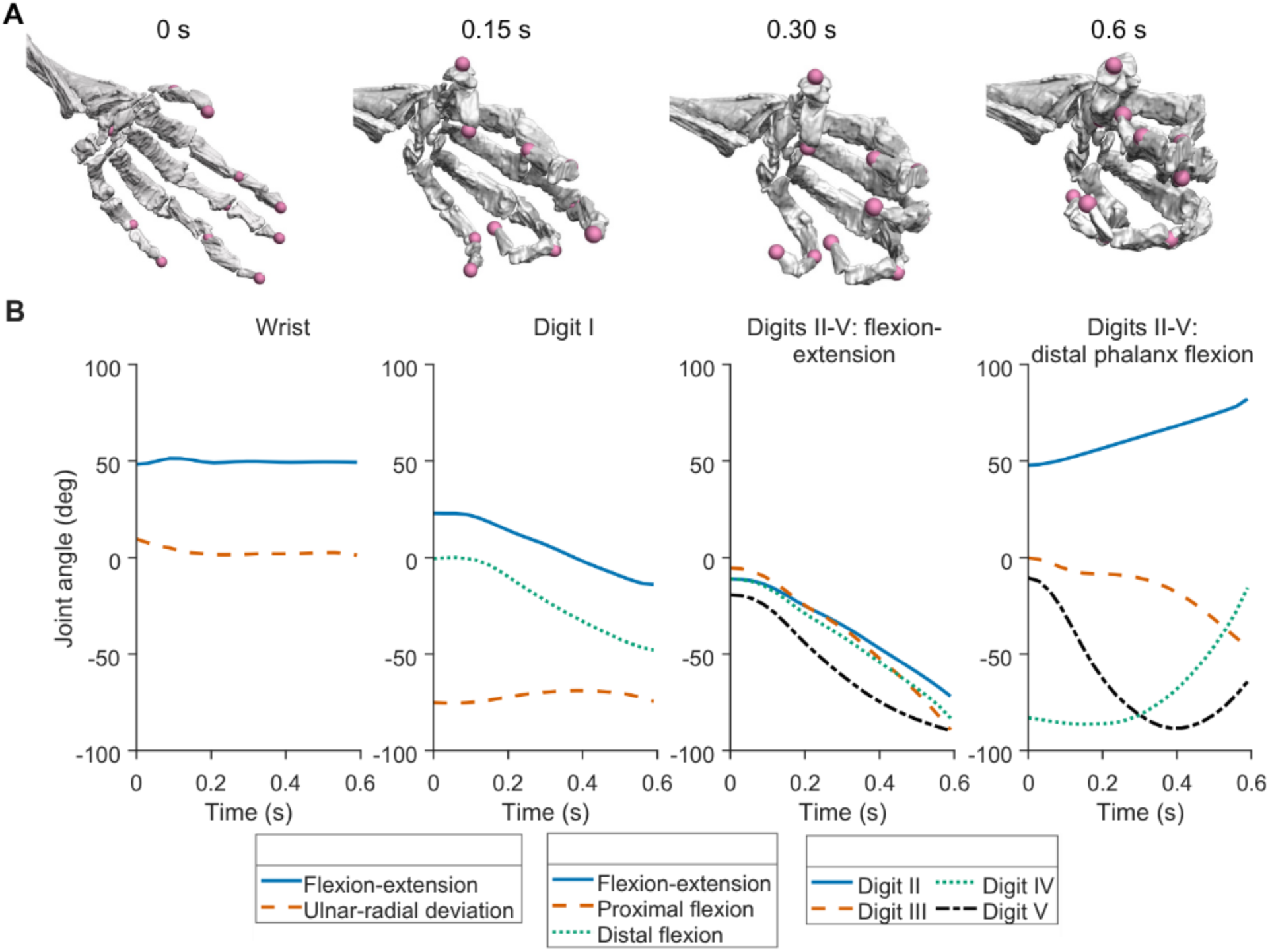
Simulated grasping movement and joint kinematics of the distal mouse forelimb. (A) Time progression of the grasping task from an initial open position at 0 s to a fully closed grasp at 0.6 s. Intermediate configurations are shown at 0.15 s and 0.30 s. The model is shown from a consistent viewpoint, focusing on the wrist and digits to highlight coordinated flexion across the fingers and digit I. Spherical markers indicate anatomical landmarks used to define the kinematic trajectories. (B) Corresponding joint-angle trajectories obtained from the simulated kinematics used to generate the grasping movement. From left to right, the plots show wrist flexion–extension and ulnar–radial deviation; digit I flexion–extension, proximal phalanx flexion, and distal phalanx flexion; flexion–extension of digits II–V; and distal phalanx flexion of digits II–V.

### Muscle-driven and Torque-driven Simulations Analysis

To evaluate agreement between torque-driven and muscle-driven solutions, marker trajectories, that is, three-dimensional positions of anatomical landmarks over time, from both simulations were compared. For each marker, the root mean square error (RMSE) was calculated to quantify discrepancies between the two approaches. For each marker and movement, the error was computed from the three–dimensional differences between corresponding marker positions (X, Y, Z) in the muscle-driven and torque-driven marker trajectories at each time point. This resulted in one RMSE value per marker for each movement, which was used to evaluate model performance across different tasks.

For validation, time-shuffled RMSE values were generated by randomly permuting the temporal order of each marker trajectory while preserving the original spatial positions, after which RMSE was recomputed using the shuffled trajectories to establish a baseline for comparison. For each marker, 500 independent random permutations were generated. A single random permutation was applied simultaneously to the X–, Y–, and Z–coordinate trajectories, preserving the spatial relationship among coordinates while disrupting their temporal correspondence. Independent permutations were generated for each marker, and the reported time-shuffled RMSE value corresponds to the mean RMSE across all 500 permutations. No fixed random seed was specified.

### EMG processing and excitation comparison

Synthesized muscle excitation profiles were extracted for comparison with experimental EMG recordings, which were obtained from the publicly available preprocessed dataset accompanying Koh et al. (2025) (Koh *et al*., 2025). Specifically, the control_centered_EMG.mat file from the Figshare repository (https://doi.org/10.6084/m9.figshare.29965298.v2) was used. The dataset contained recordings of forelimb muscle activity during naturalistic climbing in mice, including EMG from the extensor carpi radialis and palmaris longus muscles. In the original study, EMG recordings were downsampled to 1 kHz, high-pass filtered at 250 Hz, rectified, and convolved with a modified Gaussian filter kernel before being z-score normalized within each recording session (Koh *et al*., 2025).

Processed EMG recordings from the extensor carpi radialis and palmaris longus were used for comparison with the model-predicted muscle excitations because they were the only muscles shared between the experimental dataset and the distal forelimb musculoskeletal model. A reference reach from the processed dataset was used to identify the five most similar EMG profiles using the Pearson correlation-based similarity. These reaches were temporally aligned, and the mean ± standard deviation EMG profiles were computed for comparison with the simulated grasping-with-supination task generated using OpenSim Moco. Model–predicted muscle excitations were extracted from the Moco solution output for the simulated task (Dembia *et al*., 2020).

Because the model contains separate extensor carpi radialis longus and brevis muscle actuators, simulated extensor carpi radialis excitation was calculated as the unweighted arithmetic mean of the longus and brevis excitation profiles. Simulated palmaris longus excitation was extracted directly from the corresponding muscle actuator. The simulated excitation profiles were time-normalized and linearly interpolated to the same number of samples as the processed EMG traces, enabling qualitative comparison at corresponding normalized time points without modifying the experimental EMG recordings.

Experimental EMG traces were independently min-max normalized to the range of 0–1 for comparison with the model-predicted muscle excitations, which are also between 0 (no muscle activation) and 1 (maximum muscle activation). The comparison was used to evaluate the temporal agreement between experimentally observed muscle activity and model–predicted excitation patterns during grasping–related behavior.

## Results

### Marker trajectory comparison

Torque-driven and muscle-driven marker trajectories were compared to assess agreement between simulation approaches. Representative markers, including the wrist, the tip of the distal phalanx of digit I, and the tip of the distal phalanx of digit V, were selected to capture wrist and distal–digit motion during a simulated grasping task. For each marker, three-dimensional position components (X, Y, Z) were plotted over time, and trajectories from both simulation methods were overlaid to enable direct comparison of kinematic profiles.

As shown in Figure 4AB, muscle-driven and torque-driven simulations produced closely overlapping marker trajectories across all markers and spatial components. Agreement between approaches was maintained throughout the movement, with only minimal differences observed between corresponding trajectories. The distal markers exhibited larger positional changes than the wrist marker; however, the correspondence between simulation methods remained consistent across all cases. These results indicate that the muscle-driven formulation tracked the reference kinematics generated by the torque-driven simulation for the representative markers analyzed.

**Fig. 4.**
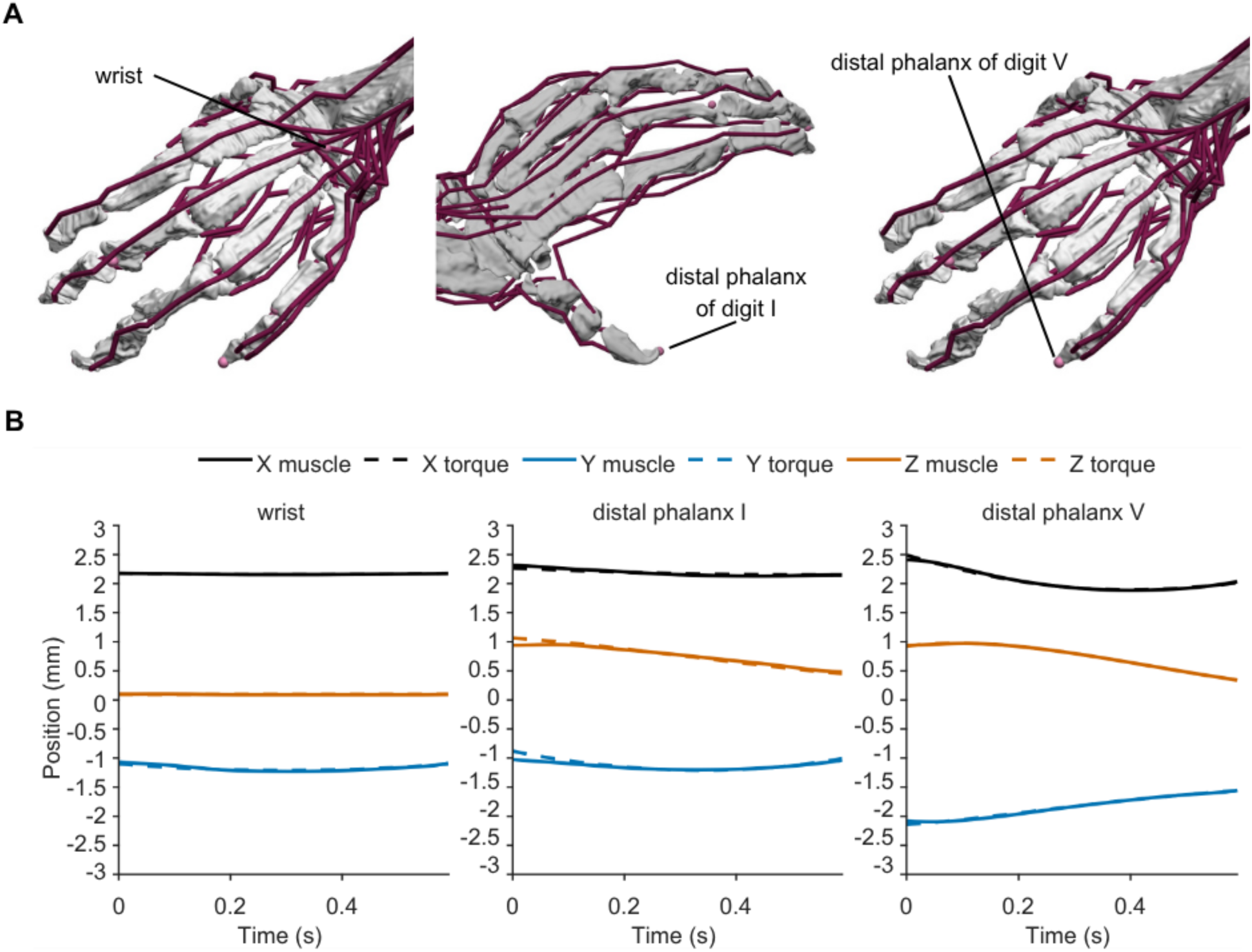
Marker trajectory comparison showing muscle-driven trajectories relative to reference trajectories generated from the torque-driven solution. Panel A illustrates the anatomical locations of the representative markers used for analysis: wrist, distal phalanx of digit I, and distal phalanx of digit V. Panel B shows the corresponding three-dimensional marker trajectories during a simulated grasping task. Position components (x, y, z) are plotted in millimeters (mm) as a function of time (s) for both muscle–driven and torque–driven simulations. Trajectories from both approaches are overlaid to enable direct comparison of marker trajectories across marker and spatial dimensions.

### RMSE for muscle-driven simulations

RMSE values varied across both movement types and marker locations in the musculoskeletal simulations (Fig. 5A). RMSE values ranged from approximately 0.017 mm to 0.131 mm across all markers–movement combinations. The largest RMSE was observed for the distal phalanx V marker during grasping with supination (0.131 mm), whereas the smallest errors were observed for the wrist marker during grasping and digit I flexion (0.017 mm). During wrist flexion, the distal phalanx 1 marker exhibited the largest RMSE values among the three markers. Grasping with supination produced the largest overall errors, whereas grasping and digit I flexion showed the lowest. Distal markers generally exhibited larger RMSE values than the wrist marker.

**Fig. 5.**
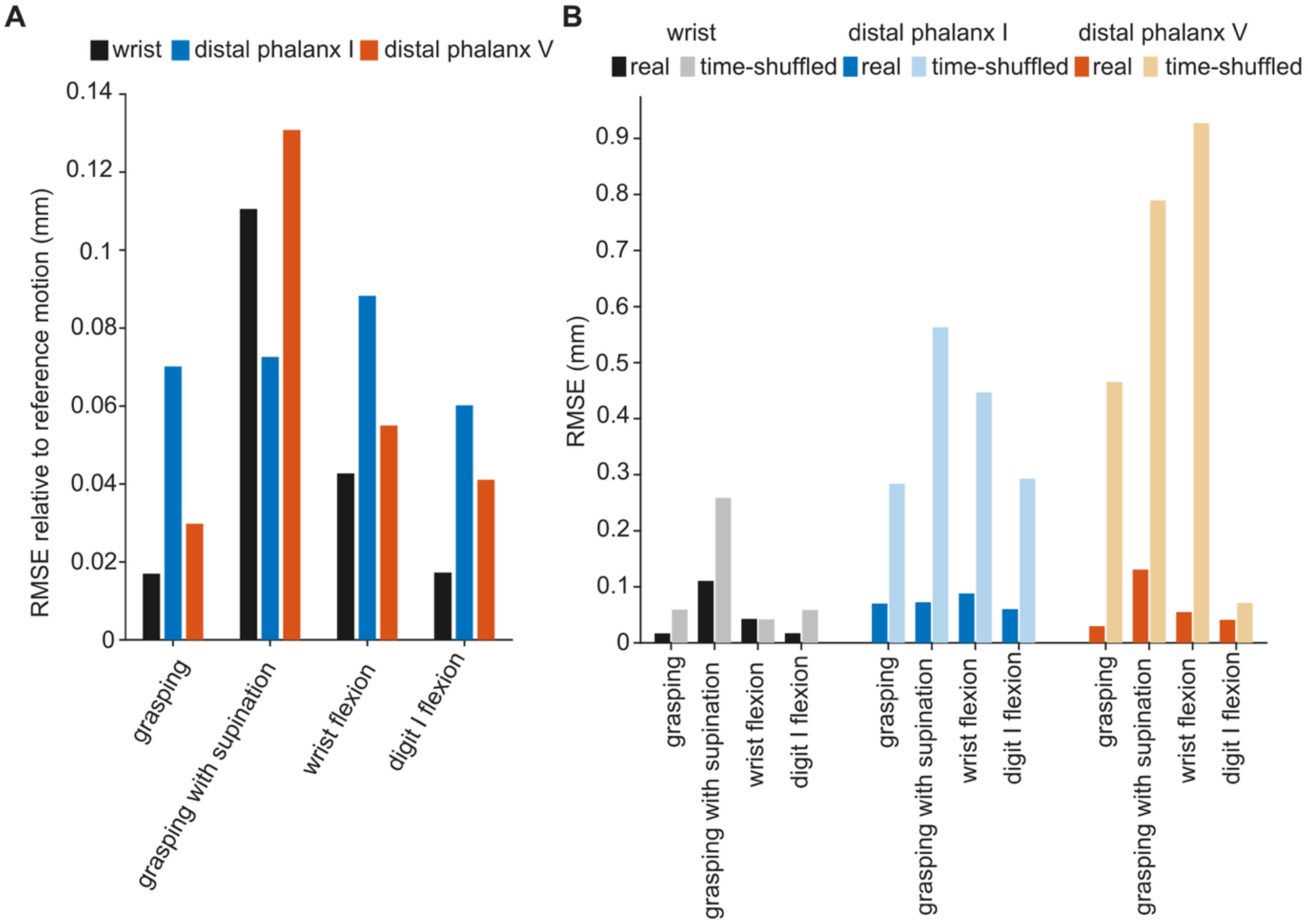
Root mean square error (RMSE) comparison across movements and with a temporal shuffling comparison. (A) RMSE of marker trajectories from the muscle–driven simulations relative to the corresponding reference trajectories generated from the torque–driven simulations for the wrist, distal phalanx of digit I, and distal phalanx of digit V across four movements: grasping, grasping with supination, wrist flexion, and digit I flexion. (B) Comparison of RMSE values obtained from the unshuffled and time–shuffled trajectory pairings for each marker across the four movements. Marker groups are arranged from left to right as the wrist, distal phalanx of digit I, and distal phalanx of digit V. Within each marker group, the darker bars represent the unshuffled pairing, labeled “real” in the panel, and the lighter bars represent the corresponding time–shuffled pairings. All RMSE values are reported in millimeters.

The time–shuffled RMSE values (Fig. 5B) were consistently higher than the corresponding unshuffled RMSE values. The distal phalanx V marker exhibited the largest increase in RMSE under time–shuffled conditions, with the greatest increases observed during wrist flexion and grasping with supination, while a comparatively smaller effect was observed during digit I flexion. In contrast, the distal phalanx I marker showed consistently large increases in RMSE across all movements when time-shuffled, showing that temporal shuffling increased its RMSE across every task. The wrist marker displayed comparatively smaller increases in RMSE across movements, with the largest difference occurring during grasping with supination.

### Muscle excitation patterns

Simulated muscle excitation patterns during the grasping task were analyzed to describe temporal variation along selected intrinsic and extrinsic muscles. As shown in Figure 6, simulated muscle excitations generated by OpenSim Moco are reported on their native scale (0–1). The panel represents six selected excitation profiles spanning intrinsic and extrinsic musculature.

**Fig. 6.**
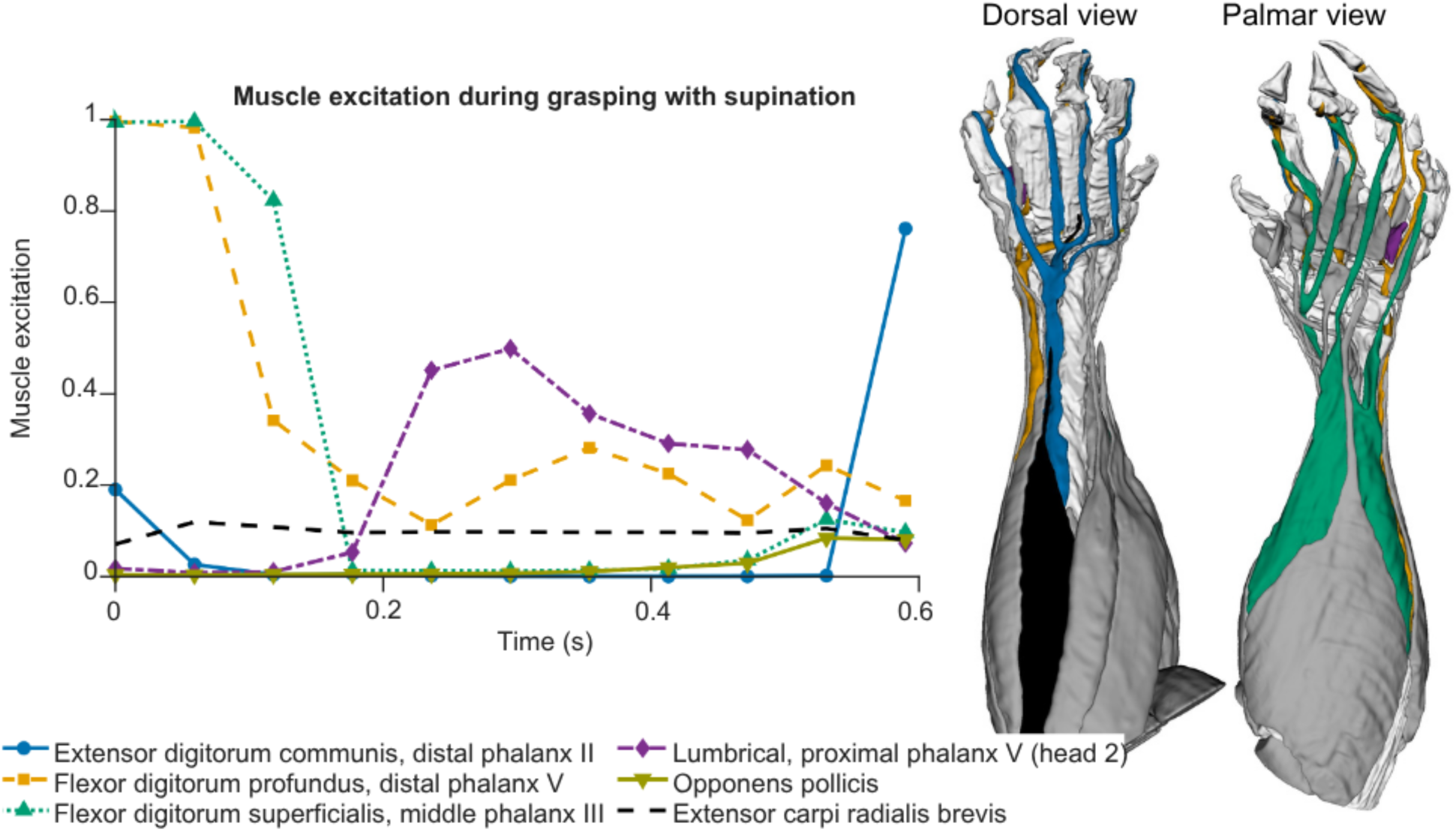
Selected muscle excitation patterns during simulated grasping with supination. Simulated excitation profiles for six selected intrinsic and extrinsic forelimb muscles during the grasping movement. Muscle excitation signals generated by the muscle-driven simulation are shown on their native normalized scale (0–1) as a function of time (0– 0.6 s). The selected muscles are extensor digitorum communis, distal phalanx II; flexor digitorum profundus radial head, distal phalanx V; flexor digitorum superficialis, middle phalanx III; lumbrical, proximal phalanx V (head 2); opponens pollicis; and extensor carpi radialis brevis. Dorsal and palmar anatomical views show the muscle groups associated with the selected excitation trajectories, using corresponding colors. Because the available anatomical muscle geometries were not subdivided into all digit-specific compartments, whereas the OpenSim model represented these compartments as separate muscle actuators, the highlighted structures provide anatomical context rather than a one-to-one visualization of each modeled actuator.

Across the six displayed profiles, excitation varied in magnitude and timing among individual muscles. The extensor digitorum communis remained near zero through most of the movement before increasing sharply near the end. The flexor digitorum profundus radial head and flexor digitorum superficialis began near maximal excitation and decreased during the first portion of the movement, although the flexor digitorum profundus radial head retained variable excitation later in the task. The lumbrical increased during the middle portion of the movement and then gradually declined, whereas the opponens pollicis remained low until a small increase near the end. Extensor carpi radialis brevis remained comparatively low and stable throughout the movement. Overall, these selected profiles demonstrate muscle-specific temporal variation but are insufficient for establishing a group-level difference between intrinsic and extrinsic musculature (Fig. 6).

### EMG excitation comparison

Simulated muscle excitation profiles generated during the grasping with supination task were compared with published EMG activity recorded during naturalistic climbing for extensor carpi radialis and palmaris longus, the only muscles shared between the experimental dataset and the distal forelimb musculoskeletal model. The experimental and simulated profiles were compared across normalized movement time, revealing both qualitative similarities and differences in their temporal behavior (Fig. 6).

Extensor carpi radialis showed a similar overall increasing trend in the experimental EMG and simulated muscle excitation profiles (Fig. 7A). The experimental EMG increased gradually across most of the movement, whereas the simulated excitation increased more slowly initially and more steeply during the latter portion of the movement. Palmaris longus showed less temporal correspondence between the experimental and simulated profiles (Fig. 7B). Experimental EMG activity was highest near the beginning of the movement and progressively decreased, whereas the simulated excitation increased to its maximum early in the movement, remained elevated through approximately the first third of the sequence, and subsequently decreased. Although both palmaris longus profiles approached their lowest activity near the end of the movement, their peak timing and early–to–middle temporal behavior differed. Overall, extensor carpi radialis showed greater qualitative temporal correspondence between experimental EMG and simulated excitation than palmaris longus.

**Fig. 7.**
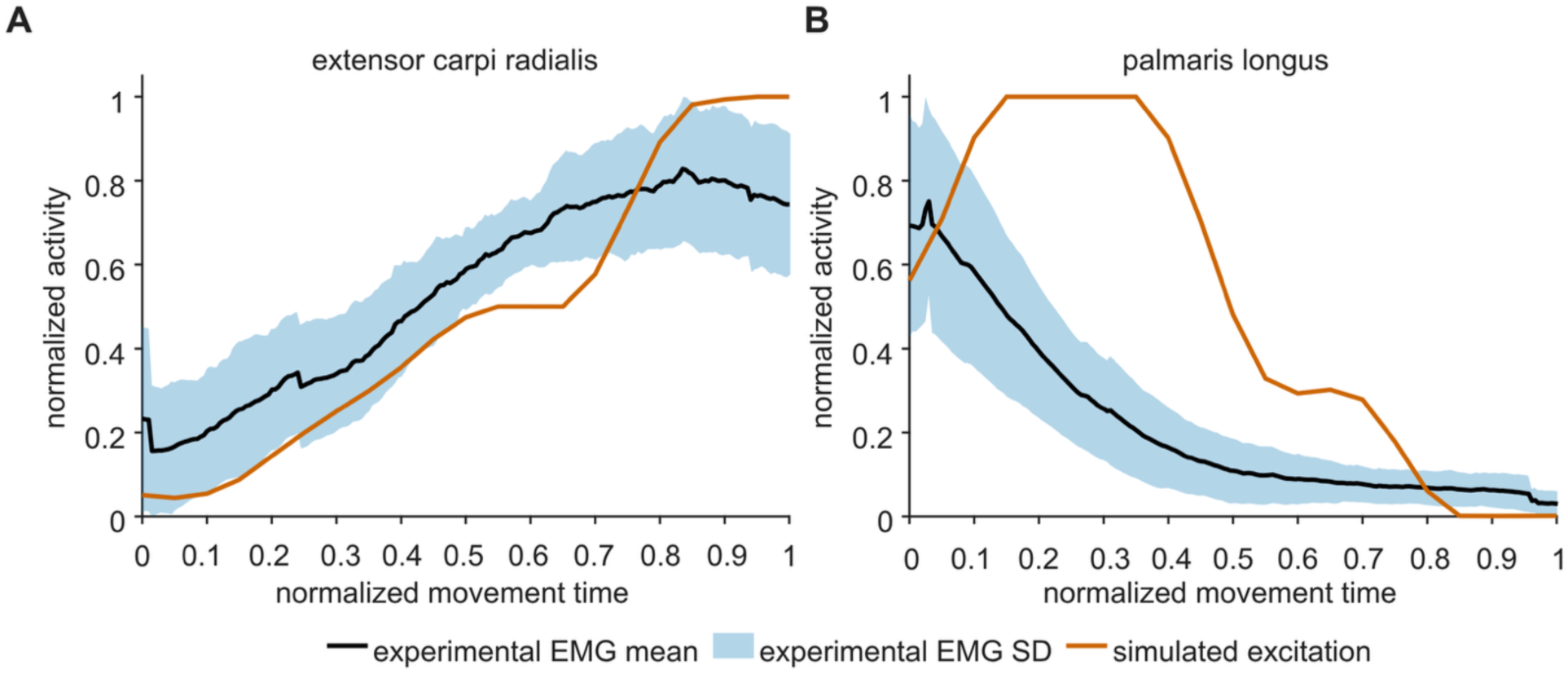
Comparison of published experimental EMG activity recorded during naturalistic climbing and simulated muscle excitation during grasping with supination. (A) Extensor carpi radialis and (B) palmaris longus. Experimental EMG recordings are shown as the mean ± standard deviation across selected trials (i.e., grasping with supination movements), while the corresponding simulated muscle excitation profile is overlaid in each panel for qualitative comparison. Both signals are displayed as independently normalized activity over normalized movement time.

## Discussion

This study developed a light-sheet-derived musculoskeletal model of the mouse distal forelimb. The muscle-driven simulations tracked digit-level marker trajectories generated by torque-driven simulations using the same model. In addition to kinematic agreement, the model showed strong correspondence between muscle-driven and torque-driven simulations, as demonstrated by the low RMSE values observed across all simulated movements, indicating consistency across simulation approaches. The temporal-shuffling comparison further showed that the low RMSE values depended on the correct temporal ordering of the reference trajectories, because shuffled pairings produced higher RMSE values. These findings indicate that the model can generate muscle-driven solutions that track prescribed distal forelimb movements, demonstrating its utility as a framework for muscle-based simulation of distal forelimb movement (Clancy *et al*., 2023). Although the representative analyses presented here focused on the grasping task, consistent tracking performance across all simulated tasks further supports the ability of the model to track internally generated reference trajectories under the tested conditions.

The simulated muscle excitation showed variation in excitation magnitude and timing among individual muscles during the grasping task (Fig. 6). The selected extrinsic and intrinsic muscles exhibited a range of excitation patterns throughout the simulated movement, including elevated initial excitation, sustained activity, decreasing excitation, and mid– or late–movement peaks. No clear group–level difference in temporal behavior was apparent between the intrinsic and extrinsic muscle groups. Accordingly, the current results do not provide sufficient evidence to attribute different functional coordination strategies to the two muscle groups. Together, these patterns demonstrate variation among the selected muscles but do not establish distinct organizational strategies for the intrinsic and extrinsic muscle groups. Previous experimental studies have shown that grasp–related muscle coordination can be modulated by grasp type, loading, wrist posture, and object–transport demands (Winges *et al*., 2007; Johnston, Bobich and Santello, 2010; Tagliabue *et al*., 2015; Geed and van Kan, 2017). Additional experimental muscle–activity measurements and quantitative group–level comparisons would be required to determine whether intrinsic and extrinsic muscles exhibit systematically different coordination patterns during mouse grasping.

The muscle-specific excitation patterns generated by the model support the value of representing distal forelimb anatomy when studying fine motor control (Milliken, Plautz and Nudo, 2013; Sobinov and Bensmaia, 2021). Existing musculoskeletal models of the mouse forelimb vary in their representation of distal skeletal structures, musculature, and tendon architecture (Ramalingasetty *et al*., 2021; Gilmer *et al*., 2024; DeWolf *et al*., 2025). In these models, distal forelimb anatomy remains simplified, with incomplete representation of the wrist, digits, and intrinsic musculature (Gilmer *et al*., 2024; DeWolf *et al*., 2025). As a result, these frameworks provide less anatomical resolution for investigating digit-level kinematics or the interaction between intrinsic and extrinsic muscles (McFarland *et al*., 2023; Gilmer *et al*., 2024). In contrast, the model developed in this study incorporates detailed anatomical structures of the distal forelimb, including all digit bones, metacarpals, phalanges, and carpal connections, as well as tendon branching and muscle paths derived from the reconstructed anatomy, including paths that pass through the carpal tunnel (DeLaurier *et al*., 2008; Watson *et al*., 2009). The inclusion of intrinsic musculature, including lumbricals, and dorsal interossei, allows the model to represent digit–level kinematics (Rath, 2011; Kooi *et al*., 2024) and provides a framework for generating testable hypotheses about distal forelimb biomechanics and muscle coordination.

Additionally, simulated muscle excitations qualitatively resembled experimentally recorded EMG activity for a grasping with supination task (Fig. 7; Koh *et al*., 2025). Extensor carpi radialis showed a broadly similar increasing trend in the experimental and simulated profiles, whereas palmaris longus showed a broadly similar decreasing trend. The palmaris longus profiles had more differences in peak timing and early-to-middle temporal behavior. In future work, we aim to evaluate the predicted muscle-excitations with EMG in other distal forelimb tasks.

Despite the strengths of the model, several limitations should be acknowledged. Muscle-specific parameters, including maximum isometric force, pennation angle, and tendon characteristics, were not directly measured for the distal forelimb and were instead estimated from previously published musculoskeletal data and scaled using the reconstructed anatomy. In addition, the maximum isometric force assigned to each muscle actuator was multiplied by a uniform factor of 1000 before simulation. Because this factor was not derived from direct physiological measurements, absolute muscle-force and excitation magnitudes should be interpreted as model-dependent estimates. The modeled intrinsic musculature included the lumbricals serving digits II–V, dorsal interossei represented through multiple muscle paths, abductor pollicis brevis, flexor pollicis brevis, opponens pollicis, flexor digiti minimi, and opponens digiti minimi. However, the abductor pollicis, abductor digiti minimi, and three palmar interossei were not included because they could not be reliably identified during three–dimensional reconstruction (Ziermann *et al*., 2021; Chen *et al*., 2026). The musculotendinous representation of the lumbricals was also simplified because their origins from the flexor digitorum profundus radial head tendons and their distinct tendon boundaries could not be reliably resolved (Chen *et al*., 2026). Consequently, the lumbricals paths were represented using bone–referenced attachments points and zero tendon slack lengths. Furthermore, because the anatomical reconstruction was derived from a single adult female C57BL/6 mouse, the model may not capture anatomical variation associated with sex, age, strain, or individual differences. The joint ranges of motion, muscle moment arms, and predicted muscle behavior were not evaluated against direct, task-matched experimental measurements. The torque-driven reference trajectories were generated using the same model subsequently evaluated through muscle–driven tracking; therefore, the RMSE analysis represents internal verification rather than independent validation. Similarly, the temporal–shuffling comparison shows that trajectory correspondence depends on temporal ordering but does not validate physiological movement. The EMG comparison was limited to two shared muscles and the experimental and simulated grasping with supination behaviors differed as the former was extracted from a climbing motion. These limitations may influence the generalizability and physiological interpretation of the predicted behavior and represent opportunities for future refinement and experimental evaluation of the model.

Despite these limitations, the model provides a framework for simulating distal forelimb movement at the digit level and could support future analyses of how neurological disorders and motor impairments affect distal forelimb function after further experimental evaluation (Young *et al*., 2026). By extending biomechanical modeling beyond the proximal limb, this approach may support future investigation of conditions such as stroke (Li *et al*., 2025), Parkinsonian motor impairment (Jasimi Zindashti *et al*., 2024), and other neurological disorders with greater distal anatomical resolution than previous proximal–limb models (Ramalingasetty *et al*., 2021; Gilmer *et al*., 2024; DeWolf *et al*., 2025). In this context, the model supports future work aimed at generating hypotheses about mechanisms of motor deficits and evaluating movement-related interventions in a more anatomically detailed and functionally relevant framework (Lorenz, Su and Skjæret-Maroni, 2024). The model will be made freely available as an open–source resource to support reuse and further evaluation by the research community.

This study presents an anatomically detailed biomechanical model of the mouse distal forelimb that incorporates digit-level skeletal anatomy, intrinsic musculature, and reconstruction-based tendon routing. The muscle-driven simulations tracked reference marker trajectories from torque-driven simulations across multiple tasks and demonstrated consistent agreement between reference marker trajectories and muscle-driven simulations. In addition, comparison between experimentally recorded EMG activity and model-predicted muscle excitation profiles identified a broad temporal similarity for the extensor carpi radialis and palmaris longus muscles, with more differences for the latter muscle. Together, these findings establish the model as a framework for generating testable hypotheses about distal forelimb biomechanics and muscle coordination at the digit level. The integration of this distal forelimb model with future full-limb biomechanical frameworks and task-matched experimental measurements may further support investigation of neurological disorders, motor deficits, and neurorehabilitation interventions in mice.

## Data availability

The OpenSim model and scripting tools used in the study and the anatomical imaging and reconstruction files are available at https://simtk.org/projects/mousehand

## Supplemental material

Supplementary information contains complete modeled muscle attachment coordinates (Table S1), muscle-tendon parameters (Table S2), marker-trajectory comparisons for three additional simulated movements (Fig. S1), and corresponding excitation profiles (Figs S2–S4), and a supplementary video showing the simulated grasping-with-supination movement.

## Grants

This work was supported by the National Institute on Drug Abuse (NIDA) of the National Institutes of Health (NIH) under Award Number R01DA060790.

## Disclosures

No conflicts of interest, financial or otherwise, are declared by the authors.

## Author contributions

N.L.S., J.I.G., G.C., D.H., and M.A.B. conceived and designed research; N.L.S., J.I.G., and G.C. performed experiments; N.L.S., J.I.G. and M.A.B. analyzed data; N.L.S., J.I.G., and M.A.B. interpreted results of experiments; N.L.S. prepared figures; N.L.S. and M.A.B. drafted manuscript; N.L.S., J.I.G., G.C., D.H., and M.A.B. edited and revised manuscript; all authors approved final version of manuscript.

## Notes

https://simtk.org/projects/mousehand

### Competing Interest Statement

The authors have declared no competing interest.

### Summary of Updates

The author order was incorrect in the first version of manuscript.

