## Supplemental for "A Physiologically Detailed Biomechanical Model of the Mouse Distal Forelimb for Simulation of Fine Motor Control"

### SUPPLEMENTARY MATERIAL

**Table S1.** *Complete muscle origins and insertions.* Origin and insertion parent bodies and corresponding three–dimensional coordinates are provided for all muscles included in the mouse forelimb musculoskeletal model. Coordinates are reported as (x, y, z) in meters and expressed in the local coordinate system of the associated parent body. Separate rows represent digit–specific muscle branches or distinct muscle heads.

| Muscle name | Origin parent-body | Origin coordinates, x, y, z (m) | Insertion parent-body | Insertion coordinates, x, y, z (m) |
|---|---|---|---|---|
| Flexor carpi ulnaris, radial head | Ulna | (-6.01E-04, -4.32E-04, -7.47E-04) | Ulnar carpal | (3.52E-04, -2.94E-04, 1.59E-04) |
| Flexor carpi ulnaris, carpal-4 head | Humerus | (-5.02E-04, -6.71E-04, -1.83E-04) | Carpal IV | (4.65E-04, -3.04E-03, 3.05E-04) |
| Flexor digitorum superficialis, middle phalanx-V | Ulna | (-2.91E-04, -1.10E-03, -8.00E-04) | Middle phalanx V | (7.88E-04, -3.70E-03, 1.26E-04) |
| Flexor digitorum superficialis, middle phalanx-IV | Ulna | (-2.91E-04, -1.10E-03, -8.00E-04) | Middle phalanx IV | (9.86E-04, -3.84E-03, 4.53E-04) |
| Flexor digitorum superficialis, middle phalanx-III | Humerus | (-3.99E-04, -7.36E-04, -4.29E-04) | Middle phalanx III | (1.04E-03, -3.83E-03, 7.33E-04) |
| Flexor digitorum superficialis, middle phalanx-II | Humerus | (-3.99E-04, -7.35E-04, -4.26E-04) | Middle phalanx II | (1.02E-03, -3.61E-03, 9.95E-04) |
| Flexor digitorum profundus, distal phalanx V | Ulna | (-1.42E-04, -1.24E-03, -7.97E-04) | Distal phalanx V | (7.62E-04, -3.93E-03, 2.35E-04) |
| Flexor digitorum profundus, distal phalanx IV | Ulna | (-1.41E-04, -1.24E-03, -8.03E-04) | Distal phalanx IV | (9.84E-04, -4.19E-03, 7.07E-04) |
| Flexor digitorum profundus, distal phalanx III | Ulna | (-1.43E-04, -1.24E-03, -7.96E-04) | Distal phalanx III | (9.95E-04, -3.98E-03, 1.06E-03) |
| Flexor digitorum profundus, distal phalanx II | Ulna | (-1.45E-04, -1.24E-03, -7.97E-04) | Distal phalanx II | (8.34E-04, -3.76E-03, 1.14E-03) |
| Extensor carpi radialis longus | Humerus | (-1.52E-04, -4.28E-04, -9.20E-04) | Metacarpal II | (6.72E-04, -2.90E-03, 5.55E-04) |
| Brachioradialis | Humerus | (-9.90E-05, -4.24E-04, -7.92E-04) | Radius | (5.30E-04, -1.99E-03, 6.50E-05) |
| Extensor digitorum communis, distal phalanx II | Humerus | (-2.75E-04, -7.13E-04, -9.42E-04) | Distal phalanx II | (8.95E-04, -3.70E-03, 1.22E-03) |
| Extensor digitorum communis, distal phalanx III | Humerus | (-2.83E-04, -7.08E-04, -9.46E-04) | Distal phalanx III | (1.08E-03, -4.04E-03, 9.85E-04) |
| Extensor digitorum communis, distal phalanx IV | Humerus | (-2.83E-04, -7.05E-04, -9.45E-04) | Distal phalanx IV | (1.06E-03, -4.15E-03, 6.96E-04) |
| Extensor digitorum communis, distal phalanx V | Humerus | (-2.83E-04, -7.07E-04, -9.47E-04) | Distal phalanx V | (8.40E-04, -3.91E-03, 2.21E-04) |
| Flexor pollicis longus | Radius | (1.00E-05, -1.09E-03, -5.50E-04) | Distal phalanx I | (7.38E-04, -3.10E-03, 9.63E-04) |
| Extensor carpi ulnaris | Humerus | (-1.25E-04, -6.58E-04, -1.03E-03) | Metacarpal V | (4.95E-04, -3.03E-03, 1.15E-04) |
| Extensor carpi radialis brevis | Humerus | (-1.65E-04, -4.48E-04, -1.03E-03) | Metacarpal III | (6.44E-04, -2.92E-03, 4.51E-04) |
| Extensor digiti minimi | Humerus | (-1.46E-04, -5.71E-04, -1.10E-03) | Distal phalanx V | (8.12E-04, -3.69E-03, 1.93E-04) |
| Abductor pollicis longus | Radius | (2.50E-05, -1.04E-03, -5.39E-04) | Carpal I | (4.87E-04, -2.89E-03, 8.18E-04) |
| Extensor pollicis longus | Ulna | (2.30E-04, -1.58E-03, -3.33E-04) | Distal phalanx I | (8.34E-04, -3.10E-03, 9.69E-04) |

| Muscle name | Origin parent-body | Origin coordinates, x, y, z (m) | Insertion parent-body | Insertion coordinates, x, y, z (m) |
|---|---|---|---|---|
| Pronator teres | Humerus | (-3.56E-04, -5.13E-04, -2.60E-04) | Radius | (3.30E-04, -1.55E-03, -2.77E-04) |
| Palmaris longus | Humerus | (-2.30E-04, -7.24E-04, -3.83E-04) | Carpal III | (4.83E-04, -3.00E-03, 4.50E-04) |
| Flexor carpi radialis | Humerus | (-3.49E-04, -6.76E-04, -1.56E-04) | Carpal II | (4.80E-04, -2.94E-03, 6.78E-04) |
| Pronator quadratus | Ulna | (4.12E-04, -2.51E-03, 1.15E-04) | Radius | (4.55E-04, -2.45E-03, 1.96E-04) |
| Supinator | Humerus | (-6.90E-05, -6.97E-04, -8.31E-04) | Radius | (4.10E-04, -1.61E-03, -2.96E-04) |
| Lumbrical, proximal phalanx II | Metacarpal II | (6.77E-04, -3.00E-03, 6.02E-04) | Proximal phalanx II | (9.28E-04, -3.30E-03, 8.15E-04) |
| Lumbrical, proximal phalanx III | Metacarpal III | (6.84E-04, -3.06E-03, 4.96E-04) | Proximal phalanx III | (1.12E-03, -3.46E-03, 6.14E-04) |
| Lumbrical, proximal phalanx IV | Metacarpal III | (7.15E-04, -3.08E-03, 4.49E-04) | Proximal phalanx IV | (9.91E-04, -3.49E-03, 3.62E-04) |
| Lumbrical, proximal phalanx IV (head 2) | Metacarpal IV | (7.66E-04, -3.20E-03, 2.69E-04) | Proximal phalanx IV | (9.86E-04, -3.48E-03, 3.63E-04) |
| Lumbrical, proximal phalanx V | Metacarpal IV | (5.98E-04, -3.04E-03, 2.43E-04) | Proximal phalanx V | (6.78E-04, -3.43E-03, 1.76E-04) |
| Lumbrical, proximal phalanx V (head 2) | Metacarpal V | (5.23E-04, -3.13E-03, 1.66E-04) | Proximal phalanx V | (6.73E-04, -3.42E-03, 1.82E-04) |
| Opponens digiti minimi | Ulnar Carpal | (3.66E-04, -2.97E-03, 2.77E-04) | Metacarpal V | (5.74E-04, -3.17E-03, 5.30E-05) |
| Flexor digiti minimi | Ulnar Carpal | (3.40E-04, -2.96E-03, 2.26E-04) | Proximal phalanx V | (6.46E-04, -3.37E-03, 8.00E-06) |
| Opponens pollicis | Carpal I | (4.92E-04, -2.90E-03, 8.11E-04) | Metacarpal I | (5.94E-04, -2.97E-03, 9.48E-04) |
| Flexor pollicis brevis | Carpal I | (4.98E-04, -2.89E-03, 8.07E-04) | Proximal phalanx I | (7.34E-04, -3.07E-03, 8.22E-04) |
| Dorsal interossei, proximal phalanx II | Metacarpal I | (5.92E-04, -3.01E-03, 7.68E-04) | Proximal phalanx II | (8.98E-04, -3.25E-03, 7.49E-04) |
| Dorsal interossei, proximal phalanx II (head 2) | Metacarpal II | (7.32E-04, -3.03E-03, 6.63E-04) | Proximal phalanx II | (8.96E-04, -3.25E-03, 7.50E-04) |
| Dorsal interossei, proximal phalanx III | Metacarpal II | (6.93E-04, -3.02E-03, 5.80E-04) | Proximal phalanx III | (1.12E-03, -3.41E-03, 5.95E-04) |
| Dorsal interossei, proximal phalanx III (head 2) | Metacarpal III | (7.45E-04, -3.08E-03, 5.24E-04) | Proximal phalanx III | (1.12E-03, -3.41E-03, 5.90E-04) |
| Dorsal interossei, proximal phalanx III (head 3) | Metacarpal III | (7.29E-04, -3.08E-03, 4.08E-04) | Proximal phalanx III | (1.13E-03, -3.45E-03, 4.06E-04) |
| Dorsal interossei, proximal phalanx III (head 4) | Metacarpal IV | (5.99E-04, -3.08E-03, 3.69E-04) | Proximal phalanx III | (1.13E-03, -3.45E-03, 4.06E-04) |
| Dorsal interossei, proximal phalanx IV | Metacarpal IV | (6.31E-04, -3.08E-03, 2.48E-04) | Proximal phalanx IV | (9.64E-04, -3.50E-03, 1.39E-04) |
| Dorsal interossei, proximal phalanx IV (head 2) | Metacarpal V | (5.72E-04, -3.15E-03, 1.87E-04) | Proximal phalanx IV | (9.65E-04, -3.50E-03, 1.36E-04) |
| Abductor pollicis brevis | Carpal I | (4.98E-04, -2.89E-03, 8.07E-04) | Proximal phalanx I | (6.27E-04, -3.04E-03, 9.05E-04) |

**Table S2.** *Muscle–tendon parameters stored in the scaled mouse forelimb musculoskeletal model.* Functional classifications and muscle–tendon parameters are provided for all muscles included in the mouse forelimb musculoskeletal model. Reported parameters include maximum isometric force, optimal fiber length, pennation angle at optimal fiber length, and tendon slack length. Distinct muscle heads and digit–specific branches are listed separately.

| Functional group | Muscle name | Base maximum isometric force (N) | Optimal fiber length (m) | Pennation angle at optimal fiber length (rad) | Tendon slack length (m) |
|---|---|---|---|---|---|
| **Wrist flexors** | Flexor carpi ulnaris, radial head | 2.18E-04 | 3.11E-03 | 0.3 | 2.93E-04 |
| | Flexor carpi ulnaris, carpal-IV head | 2.57E-04 | 2.81E-03 | 0.3 | 8.20E-04 |
| | Palmaris longus | 1.84E-03 | 8.51E-04 | 0.3 | 3.37E-03 |
| | Flexor carpi radialis | 2.84E-03 | 1.10E-03 | 0.3 | 2.44E-03 |
| **Wrist extensors** | Extensor carpi radialis longus | 4.71E-03 | 9.61E-04 | 0.3 | 3.42E-03 |
| | Extensor carpi ulnaris | 1.97E-03 | 8.58E-04 | 0.3 | 3.00E-03 |
| | Extensor carpi radialis brevis | 2.30E-03 | 9.46E-04 | 0.3 | 3.58E-03 |
| **Elbow flexor** | Brachioradialis | 3.72E-03 | 9.32E-04 | 0.3 | 1.47E-03 |
| **Forearm rotators** | Pronator teres | 3.68E-03 | 1.48E-03 | 0.3 | 0.00E+00 |
| | Pronator quadratus | 9.49E-04 | 1.34E-04 | 0.1 | 0.00E+00 |
| | Supinator | 1.64E-03 | 1.39E-03 | 0.3 | 0.00E+00 |
| **Extrinsic digit flexors** | Flexor digitorum superficialis, middle phalanx-V | 2.87E-04 | 8.70E-04 | 0.3 | 3.29E-03 |
| | Flexor digitorum superficialis, middle phalanx-IV | 2.33E-04 | 1.06E-03 | 0.3 | 3.42E-03 |
| | Flexor digitorum superficialis, middle phalanx-III | 1.98E-04 | 1.62E-03 | 0.3 | 3.03E-03 |
| | Flexor digitorum superficialis, middle phalanx-II | 1.98E-04 | 1.97E-03 | 0.3 | 2.60E-03 |
| | Flexor digitorum profundus, distal phalanx V | 2.12E-04 | 1.16E-03 | 0.2 | 2.33E-03 |
| | Flexor digitorum profundus, distal phalanx IV | 2.12E-04 | 8.31E-04 | 0.2 | 3.54E-03 |
| | Flexor digitorum profundus, distal phalanx III | 2.12E-04 | 1.13E-03 | 0.2 | 3.14E-03 |
| | Flexor digitorum profundus, distal phalanx II | 2.12E-04 | 9.95E-04 | 0.2 | 3.45E-03 |
| | Flexor pollicis longus | 3.91E-04 | 4.39E-04 | 0.1 | 2.67E-03 |
| **Extrinsic digit extensors** | Extensor digitorum communis, distal phalanx II | 4.68E-04 | 9.76E-04 | 0.2 | 4.08E-03 |
| | Extensor digitorum communis, distal phalanx III | 4.68E-04 | 1.32E-03 | 0.2 | 3.81E-03 |
| | Extensor digitorum communis, distal phalanx IV | 4.68E-04 | 1.38E-03 | 0.2 | 4.11E-03 |
| | Extensor digitorum communis, distal phalanx V | 4.68E-04 | 1.87E-03 | 0.2 | 2.94E-03 |
| | Extensor digiti minimi | 1.48E-03 | 5.12E-04 | 0.2 | 4.50E-03 |
| | Abductor pollicis longus | 2.51E-03 | 1.34E-03 | 0.3 | 1.52E-03 |
| | Extensor pollicis longus | 2.42E-04 | 1.12E-03 | 0.1 | 1.70E-03 |
| **Intrinsic hand muscles** | Lumbrical, proximal phalanx II | 5.43E-04 | 5.49E-04 | 0.1 | 0.00E+00 |
| | Lumbrical, proximal phalanx III | 5.34E-04 | 6.17E-04 | 0.1 | 0.00E+00 |
| | Lumbrical, proximal phalanx IV | 7.87E-04 | 6.63E-04 | 0.1 | 0.00E+00 |
| | Lumbrical, proximal phalanx IV (head 2) | 5.25E-04 | 5.37E-04 | 0.1 | 0.00E+00 |
| | Lumbrical, proximal phalanx V | 3.21E-04 | 2.58E-04 | 0.1 | 0.00E+00 |
| | Lumbrical, proximal phalanx V (head 2) | 2.73E-04 | 4.01E-04 | 0.1 | 0.00E+00 |
| | Opponens digiti minimi | 3.43E-04 | 4.24E-04 | 0.1 | 0.00E+00 |
| | Flexor digiti minimi | 3.36E-04 | 5.94E-04 | 0.1 | 0.00E+00 |

| Functional group | Muscle name | Base maximum isometric force (N) | Optimal fiber length (m) | Pennation angle at optimal fiber length (rad) | Tendon slack length (m) |
|---|---|---|---|---|---|
| **Intrinsic hand muscles** | Opponens pollicis | 6.20E-05 | 1.74E-04 | 0.1 | 0.00E+00 |
| | Flexor pollicis brevis | 2.56E-04 | 2.57E-04 | 0.1 | 0.00E+00 |
| | Dorsal interossei, proximal phalanx II | 1.20E-05 | 5.48E-04 | 0.1 | 0.00E+00 |
| | Dorsal interossei, proximal phalanx II (head 2) | 1.90E-05 | 4.37E-04 | 0.1 | 0.00E+00 |
| | Dorsal interossei, proximal phalanx III | 2.54E-04 | 5.93E-04 | 0.1 | 0.00E+00 |
| | Dorsal interossei, proximal phalanx III (head 2) | 2.57E-04 | 5.55E-04 | 0.1 | 0.00E+00 |
| | Dorsal interossei, proximal phalanx III (head 3) | 3.04E-04 | 5.46E-04 | 0.1 | 0.00E+00 |
| | Dorsal interossei, proximal phalanx III (head 4) | 3.55E-04 | 6.06E-04 | 0.1 | 0.00E+00 |
| | Dorsal interossei, proximal phalanx IV | 1.80E-04 | 6.43E-04 | 0.1 | 0.00E+00 |
| | Dorsal interossei, proximal phalanx IV (head 2) | 1.32E-04 | 7.27E-04 | 0.1 | 0.00E+00 |
| | Abductor pollicis brevis | 4.40E-04 | 2.04E-04 | 0.1 | 0.00E+00 |

Maximum isometric force values are the base values store in handModel_scaled.osim and were multiplied by 1000 during the muscle–driven simulations. Pennation angles are specified at optimal fiber length. Zero tendon slack–length values reproduce the values store in the scaled model. Distinct muscle heads and digit–specific branches are listed separately.

**Table S3.** *Optimization settings for OpenSim Moco simulations of distal forelimb movements.* Optimization settings used for the torque-driven and muscle-driven OpenSim Moco simulations performed for the four prescribed distal forelimb movements (grasping, grasping with supination, digit I flexion, and wrist flexion). The table summarizes the software versions, optimization framework, solver settings, convergence criteria, tracking objectives, actuator configuration, and control parameters used to generate the kinematic and muscle-driven simulations described in the Methods

| Parameter | Torque-driven simulation | Muscle-driven simulation |
|---|---|---|
| Software | OpenSim 4.5 | OpenSim 4.5 |
| OpenSim Creator | 0.7.1 | 0.7.1 |
| Python | 3.13.9 | 3.13.9 |
| Optimization framework | OpenSim Moco | OpenSim Moco |
| Solver | CasADi | CasADi |
| Transcription method | Direct collocation | Direct collocation |
| Mesh intervals | 60 | 60 |
| Convergence tolerance | $1 \times 10^{-5}$ | $1 \times 10^{-4}$ |
| Constraint tolerance | $1 \times 10^{-5}$ | $1 \times 10^{-4}$ |
| Maximum optimization iterations | 500 | 500 |

| Parameter | Torque-driven simulation | Muscle-driven simulation |
|---|---|---|
| Tracking target | Prescribed joint coordinates | Three-dimensional marker trajectories |
| Marker tracking goal | N/A | MocoMarkerTrackingGoal |
| Marker identities | N/A | Wrist, Metacarpal I, Proximal Phalanges II–V, Distal Phalanges I–V |
| Marker weights | N/A | Equal weighting ($1 \times 10^9$ per marker) |
| Control effort goal | Sum of squared coordinate-actuator controls | Sum of squared muscle excitations |
| Control effort weight | 1 | 1 |
| Actuators | Coordinate actuators | Hill-type musculotendon actuators |
| Initial coordinate values | Prescribed for each simulated movement | Prescribed for each simulated movement |
| Final coordinate values | Prescribed for each simulated movement | Prescribed for each simulated movement |
| Coordinate actuator strengths | Defined in the OpenSim model | N/A |
| Coordinate actuator control bounds | Defined in the OpenSim model | N/A |
| Muscle excitation bounds | N/A | Defined in the OpenSim model |
| Muscle control bounds | N/A | Defined in the OpenSim model |

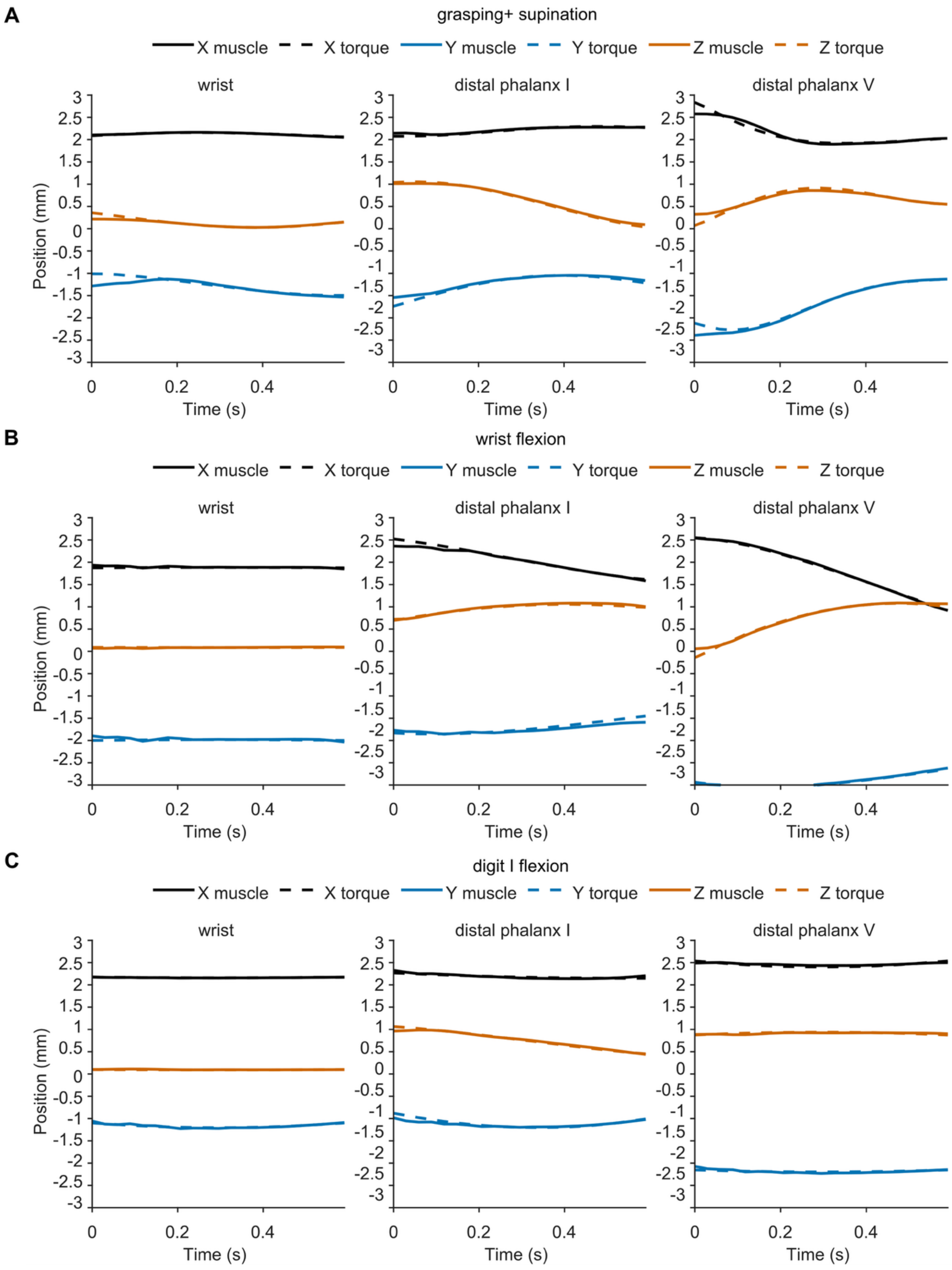


**Figure S1.** *Comparison of marker trajectories generated by muscle-driven and torque-driven*

*simulations*. Marker trajectories predicted by the muscle-driven (solid lines) and torque-driven (dashed lines) simulations for three representative markers during three validation tasks. Panel A shows trajectories during grasping with supination, panel B shows trajectories during writ flexion, and panel C shows trajectories during digit I flexion. Withing each panel, the left, middle, and right plots show the wrist, distal phalanx I, and distal phalanx V markers, respectively. Black, blue, and orange traces represent the X-, Y-, and Z-coordinate positions, respectively. The close overlap between muscle-driven and torque-driven trajectories demonstrates that the muscle-driven simulations accurately reproduced the kinematics generated by the torque-driven tracking solution across all tasks.

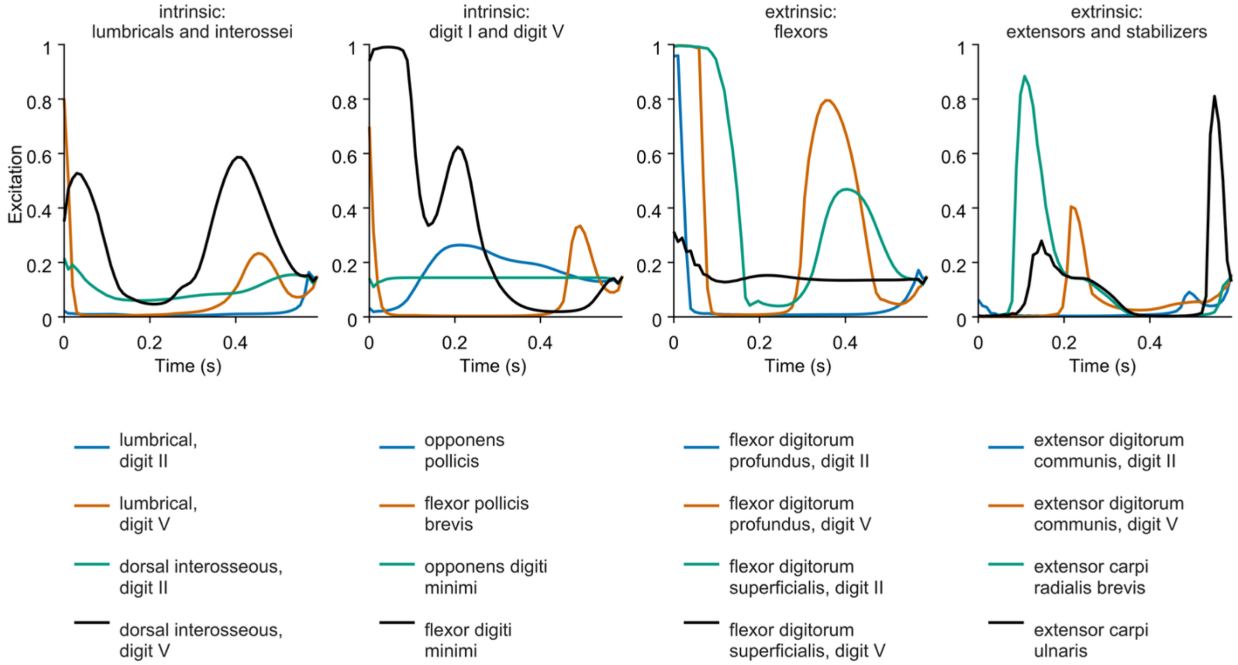


**Figure S2.** *Predicted muscle excitations generated by the muscle-driven simulation during the grasping with supination task.* From left to right, the plots show the intrinsic lumbrical and dorsal interosseous muscles of digits II and V; intrinsic muscles of digits I and V, including opponens pollicis, flexor pollicis brevis, opponens digiti minimi, and flexor digiti minimi; extrinsic digit flexors, including flexor digitorum profundus and flexor digitorum superficialis for digits II and V; and extrinsic digit extensors and wrist stabilizers, including extensor digitorum communis for digits II and V, extensor carpi radialis brevis, and extensor carpi ulnaris. Muscle excitation is normalized from 0 to 1 and plotted over the duration of the simulated movement.

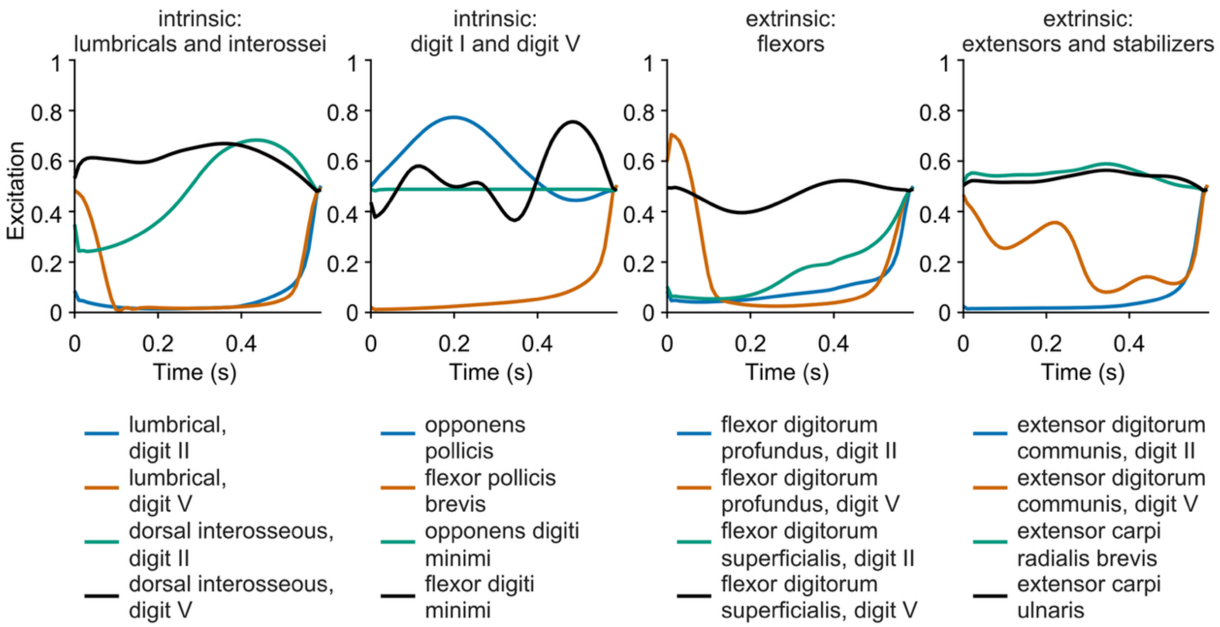


**Figure S3.** *Predicted muscle excitations generated by the muscle-driven simulation during the wrist flexion task.* From left to right, the plots show the intrinsic lumbrical and dorsal interosseous muscles of digits II and V; intrinsic muscles of digits I and V, including opponens pollicis, flexor pollicis brevis, opponens digiti minimi, and flexor digiti minimi; extrinsic digit flexors, including flexor digitorum profundus and flexor digitorum superficialis for digits II and V; and extrinsic digit extensors and wrist stabilizers, including extensor digitorum communis for digits II and V, extensor carpi radialis brevis, and extensor carpi ulnaris. Muscle excitation is normalized from 0 to 1 and plotted over the duration of the simulated movement.

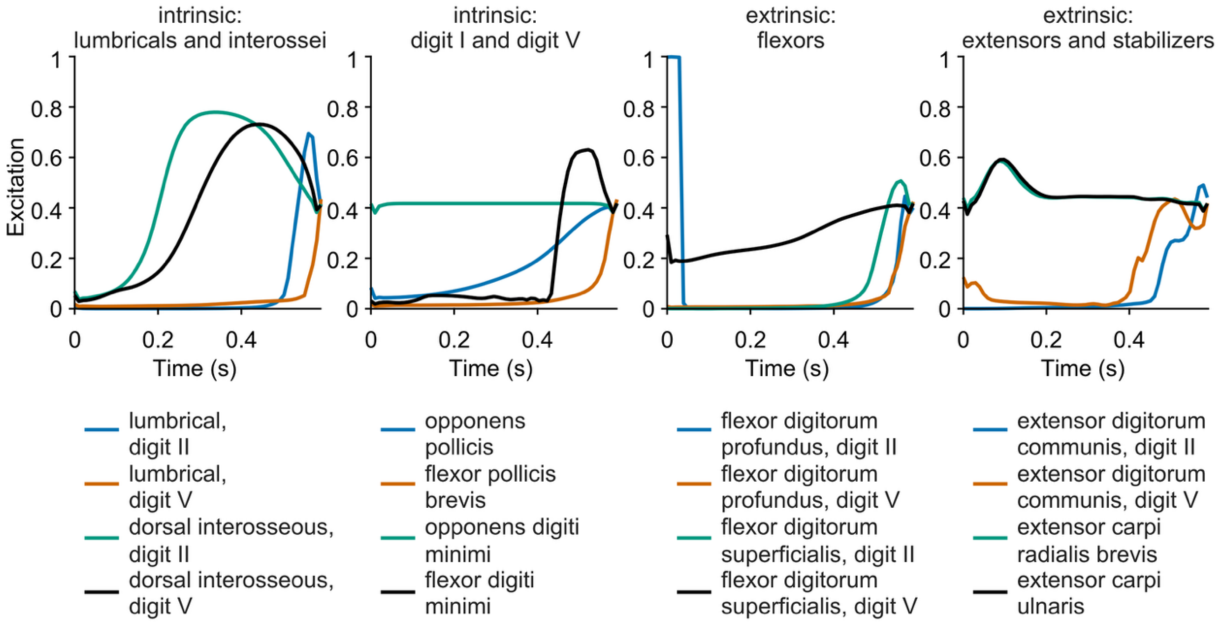


**Figure S4.** *Predicted muscle excitations generated by the muscle-driven simulation during the digit I flexion task.* From left to right, the plots show the intrinsic lumbrical and dorsal interosseous muscles of digits II and V; intrinsic muscles of digits I and V, including opponens pollicis, flexor pollicis brevis, opponens digiti minimi, and flexor digiti minimi; extrinsic digit flexors, including flexor digitorum profundus and flexor digitorum superficialis for digits II and V; and extrinsic digit extensors and wrist stabilizers, including extensor digitorum communis for digits II and V, extensor carpi radialis brevis, and extensor carpi ulnaris. Muscle excitation is normalized from 0 to 1 and plotted over the duration of the simulated movement.
